# Real-Time Embodied Experience Shapes High-Level Reasoning Under Altered Gravity

**DOI:** 10.64898/2026.03.16.712090

**Authors:** Hélène Grandchamp des Raux, Tommaso Ghilardi, Elisa Raffaella Ferrè, Ori Ossmy

## Abstract

A critical aspect of human cognition is the ability to use our knowledge about the laws of physics to make predictions about physical events. Whether this ability is based on abstract processes or is grounded in our body-environment interactions remains an open debate. We used physical reasoning under altered gravity as a model system to show that humans’ real-time embodied experience modifies their high-level physical reasoning. Specifically, we tested participants in computerised reasoning games, while disrupting their gravitational signalling using Galvanic Vestibular Stimulation (GVS). Participants failed more and had suboptimal strategies under the GVS condition compared to no-GVS in games requiring reasoning about terrestrial gravity. However, the effects of GVS were reduced when the games included reasoning about altered gravity. Our findings demonstrate how the physical experience of the body shifts high-level cognitive skill as reasoning, suggesting that humans’ mental representation of the world is grounded in adaptable physical mechanisms.

## Introduction

Physical reasoning—the intuitive ability to predict how objects and environments behave under the laws of physics—is a fundamental component of human cognition^1,2^. Even the simplest daily activities, as moving a cup away from a table’s brink or catching a ball in mid-air, require humans to harness cognitive resources to anticipate and respond to their physical environment. These daily activities are supported by the ability to use knowledge of real-world physical laws to predict environmental changes, understand which action drives a successful interaction with their environment and interpret and extract valuable insights from failed actions. Yet, despite the importance of physical reasoning for human survival, the question is still open regarding its underlying mechanisms.

A contentious debate in cognitive psychology and neuroscience has revolved around whether human reasoning is an abstract, modular process^3,4^ that relies on amodal symbols, rules and conventions^3^, or whether it is fundamentally grounded in human bodily interactions with the environment^5–8^. In other words, is physical reasoning a result of a ‘high-level’ rational, logic-based inference or is it honed and refined through ‘low-level’ embodied sensorimotor experience? This is not a trivial debate. The implications extend far beyond psychological research, informing fields such as physics education^9–12^, instructional design, artificial intelligence and robotics. This is particularly relevant given the continuing challenge of replicating human-like common sense, intuitive reasoning, and adaptive behaviour in artificial systems^13^.

Traditionally, cognitive researchers argue that physical reasoning is grounded in innate knowledge of the relationships between cues and outcomes and is largely detached from the real-time sensorimotor experiences that shape cognition^14–17^. Similarly, Bayesian models of cognition suggest that individuals use priors grounded in an intuitive knowledge of Newtonian physics, informed by past observations, to refine real-time physical predictions^15,18,19^. Under perceptual uncertainty, these priors integrate with new sensory data to forecast causal relationships and potential outcomes of actions^20–22^.

An alternative view on physical reasoning has emerged in the last decades, which argues that cognitive processes are fundamentally grounded in bodily experiences and interactions with the physical world ^3,5^. For example, humans’ prediction of the trajectory of moving objects (e.g., a ball thrown towards them) relies on their need to physically act upon this prediction (e.g., catch the ball). Moreover, the potency of physical reasoning in this context is affected by individual variations in skills^23,24^. A seasoned athlete with robust physical prowess may be more adept at foreseeing the course of a swiftly thrown ball, granting them a competitive edge in timely reactions^25,26^. This interplay between prediction and action, underpinned by physical reasoning, allows humans to efficiently navigate through dynamic environments, adapt to novel scenarios, and react with precision and agility to unpredictable situations.

Gravity’s ubiquitous presence on Earth offers a unique framework for investigating a central question in cognitive science concerning whether physical reasoning arises from high-level, abstract inferential processes or is instead grounded in, and shaped by, embodied sensorimotor experience. All living species on our planet have been continuously exposed to a constant gravitational acceleration of 9.8 m/s^2^ (1g) which has influenced the way we successfully interact with the environment^27,28^. The vestibular system in the inner ear continuously monitors the position of the head with respect to gravity. By integrating vestibular signals with visual, proprioceptive, and visceral cues, the brain creates an internal model of terrestrial gravity, which provides a fundamental prior for reasoning about motion and spatial orientation and predicting trajectories, weight, and stability of objects in everyday contexts^28^. Developmental evidence suggests that aspects of this gravity prior are innate, as events that defy expected gravitational behaviour violate the expectations of human toddlers^29^ and animals^30^.

However, it remains unclear to what extent real-time embodied experience can recalibrate this internal mental model. In other words, does immediate sensorimotor exposure to a novel gravitational environment alter high-level physical reasoning, or is this gravity model largely fixed? Addressing this question is important because it speaks to the flexibility of the human mind: if brief physical experiences can update an internal gravity model, then one of the most abstract mental skills is grounded in recent bodily states. If not, reasoning may rely on - and be limited by - a more ‘hard-wired’ prior. Here, we address this question by conducting two *pre-registered* studies, guided by a simple logic: if real-time embodied experience affects the mechanisms underlying physical reasoning, then altering a person’s gravitational cues while solving a physical reasoning task should affect their internal gravity model (and thereby their task performance).

In the first study, we tested participants under terrestrial gravity with and without perturbation of vestibular gravity cues. Healthy adult participants completed the Virtual Tools task^20^—a set of computerized reasoning games in which they had to reason about the movement of objects in a virtual 2D environment (Fig. 1a and Supplementary Video 1).

**Fig. 1.**
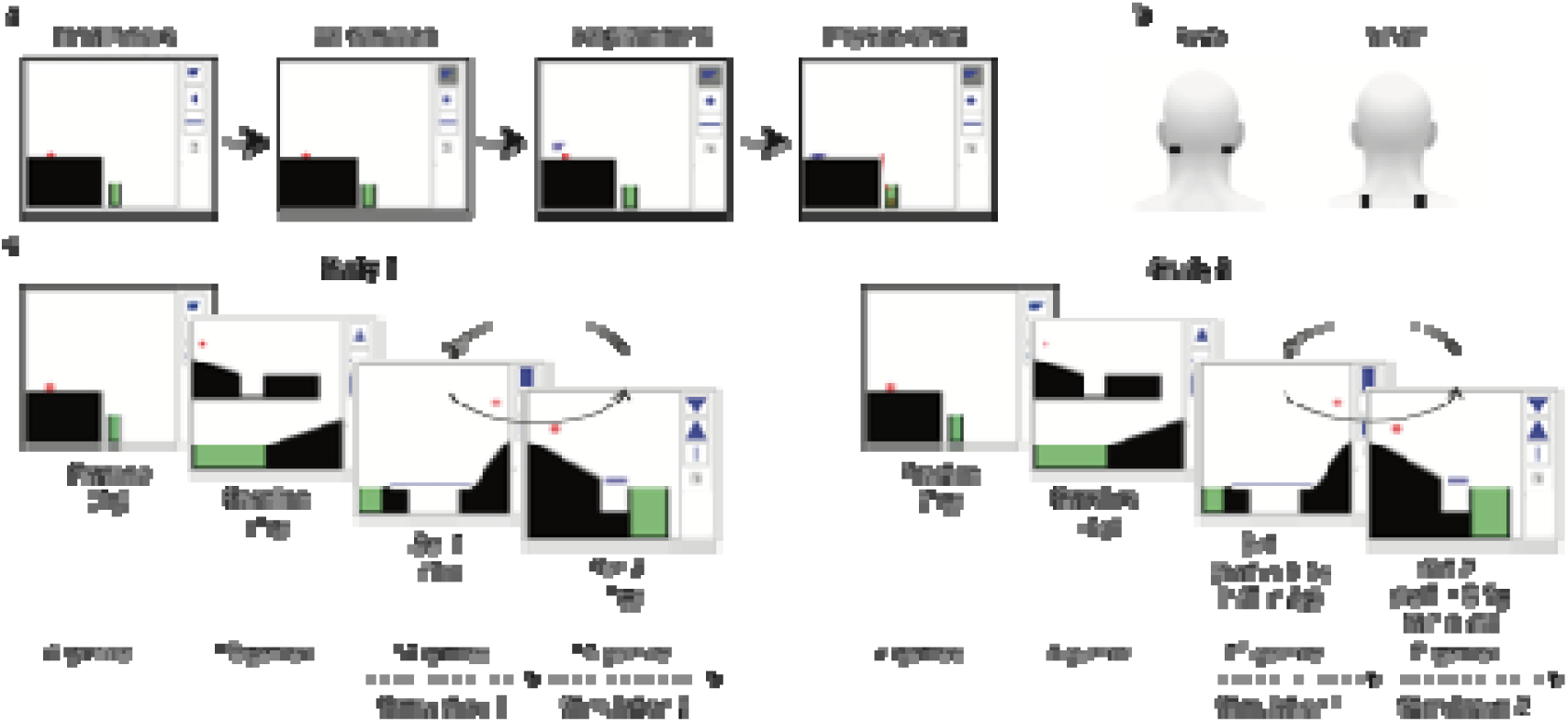
Task design. a, Virtual Tools task. The participants were required to click on one out of the three tools depicted in the upper right corner of the figure (second panel from the left) and to place it with another click on the screen to get the red ball into the green area (third panel from the left). As soon as the tool is released (panel on the right), the laws of physics/gravity are applied, and the tool interacts with the other objects. b, Galvanic Vestibular Stimulation set up with two pairs of electrodes shown on a head model. The upper electrodes deliver the Galvanic Vestibular Stimulation and the ones below are used as a Sham stimulation to control for non-vestibular specific effects. c, Procedure. After a series of practice games followed by baseline games designed under terrestrial gravity, participants played two sets of games designed under terrestrial gravity (in study 1) and hypo or hyper gravities (in study 2), with concurrent stimulation (either GVS or Sham during a given set).

While participants performed the reasoning tasks, we transiently perturbed gravitational signalling using disruptive Galvanic Vestibular Stimulation (GVS)^31^, which introduces controlled noise into vestibular afferent processing. In contrast to genuine alterations of gravitational input achieved through spaceflight and terrestrial analogues (e.g., parabolic flights, centrifugation, and head-down bed rest), this approach selectively modulates neurovestibular signalling without eliciting concomitant changes in other physiological systems, including cardiovascular, respiratory, or musculoskeletal function. Participants also completed a sham stimulation condition that reproduced comparable cutaneous sensations while leaving vestibular processing unaffected; condition order was fully counterbalanced across participants.

Participants engaged in a set of games that systematically varied in their reliance on gravity-based predictions, enabling us to test whether disrupting vestibular input selectively impairs reasoning in gravity-dependent contexts or instead produces a more general cognitive disruption. This design allowed us to isolate the specific contribution of gravitational processing to physical reasoning. The Virtual Tools task minimises motor demands and performance depends primarily on tool selection and spatial placement. We therefore focused on high-level physical reasoning measures (success rate, mean number of attempts, and mean attempt duration) alongside strategy indices, operationalised as trial-to-trial changes in tool selection and placement within each game, rather than movement kinematics. Because participants were not instructed explicitly to respond as quickly as possible, time per attempt could potentially reflect several processes, including deliberation, hesitation, task engagement, or individual speed-accuracy trade-offs, rather than better or worse reasoning. We therefore treated time per attempt as a secondary descriptive measure of attempt, rather than a direct index of performance efficiency. Better performance and more efficient strategies in the Sham condition compared to the GVS condition in gravity-dependent games would suggest that an intact real-time experience is necessary for high-level reasoning about physical dynamics. Alternatively, a similar or better performance in the GVS condition would suggest that disrupting the real-time experience of gravity does not selectively degrade reasoning performance. In our pre-registration, we hypothesised that when participants do not experience changes in gravity (sham stimulation), they would perform better in physical-reasoning tasks that are based on terrestrial gravity.

In the second study, we aimed to test whether the hindering effects of real-time embodied experience can be reduced in case of reasoning about *altered* gravity. To that end, we manipulated the gravity within the Virtual Tools games, which required reasoning about hypogravity (0.5 g) or hypergravity (2 g) while each participant experienced GVS or Sham condition. This meant that object movements were influenced by gravity at either a higher or lower acceleration than Earth’s standard gravity (9.8 m/s²). We calculated the performance and strategy measures as in study 1. Then, for each measure, we computed a Gravity-Index (GI) by calculating the ratio between GVS-performance to Sham-performance (see Methods). This GI is a direct quantification of the effects of real-time vestibular noise on physical reasoning. The use of an index, calculated in the same way in both studies, allowed us to compare the relative effect of GVS across the two experiments rather than relying on differences in raw performance between the two participant samples. For success rate, a higher GI in study 2 compared to study 1 would suggest that the effects of the real-time embodied disruption are reduced when reasoning about non-terrestrial gravity. Conversely, a lower GI would suggest that high-level reasoning depends mainly on a fixed gravity prior, and that real-time vestibular noise leaves participants’ performance unchanged, regardless of the gravity within the game. We hypothesized that participants would perform better with altered sense of gravity when the task involves non-terrestrial gravity.

## Results

### Study 1: Intact Real-Time Embodied Experience is Necessary for High-Level Reasoning

We first examined whether the altered vestibular input influenced participants’ performance in the reasoning task - defined as success rate, number of attempts to solve the games and time per attempt. High success rates and low number and time per attempt indicate good performance. Preliminary analyses showed that males and females did not perform significantly differently between the Sham and GVS conditions (male; *t*(9) = -0.58, p = .71, Cohen’s *d* = -0.22; female: *t*(33) = 0.47, p = .32, Cohen’s *d* = 0.10), so we combined all participants in subsequent analyses.

When examining all reasoning games, participants performed similarly in the Sham condition compared to the GVS condition in all performance measures (*t_success_rate_*(43) = 0.27, *p_sucess_rate_* = .39, Cohen’s *d_sucess_rate_* = 0.05; *t*_number_of_attempt_ (43) = -0.88, *p*_number_of_attempt_ = .19, Cohen’s *d*_number_of_attempt_ = -0.14; *t*_time_per_attempt_(43) = 0.32, *p*_time_per_attempt_ = .62, Cohen’s *d*_time_per_attempt_ = 0.05; See full statistics in Table 1). When grouping games based on their gravity impact level categorisation, the overall interaction between the stimulation type and the gravity impact level did not reach statistical significance across all performance measures (all x^2^ ≤ 3.55, all *p*s ≥ .17, see full statistics in Table 2 and Methods for the models’ details). However, the pairwise contrasts revealed a significant effect within the low gravity impact for time, with an average time under Sham 8.5% longer than under GVS (ratio_time_low_ = 1.08, t_time_low_(1064)= 1.99, p_time_low_ = .047) as well as a trend emerging within the gravity medium impact level for number of attempts, with a 13.4% reduction in the number of attempts under Sham compared to GVS (ratio_attempts_medium_ = 0.87, z_attempts_medium_= -1.87, p_attempts_medium_ = .061).

**Table 1.** Means, Standard Deviations and t test Statistics for Study 1 measures.

| Variable | Sham |  | GVS |  | df | t | p | Cohen's<br>d |
| --- | --- | --- | --- | --- | --- | --- | --- | --- |
|  | M | SD | M | SD |  |  |  |  |
| Success Rate (%) | 75.1 | 14.2 | 74.3 | 16.9 | 43 | 0.27 | .39 | 0.05 |
| Number of Attempts | 3.0 | 0.9 | 3.1 | 1.1 | 43 | -0.88 | .19 | -0.14 |
| Time per Attempt (s) | 6.5 | 2.3 | 6.4 | 2.6 | 43 | 0.32 | .62 | 0.05 |
| Distance (pixel) | 115.8 | 29.9 | 120.0 | 44.1 | 43 | -0.52 | .61 | -0.11 |
| Tool Switching (%) | 66.1 | 13.5 | 61.9 | 16.1 | 43 | 1.9 | .06 | 0.28 |

Paired t-tests. $p < .05$

**Table 2.** Summary of overall interactions and pairwise contrasts for Study 1.

| Model | Effect / Contrast | Estimate [95% CI] | z or t | p-value |
| --- | --- | --- | --- | --- |
| SR [Binomial Variable]<sup>a</sup> |  |  |  |  |
| | Overall interaction | $\chi^2 = 0.93$ | | .63 |
|  | Low Gravity Impact: Sham vs GVS | 0.90 | -0.28 | .78 |
|  | Medium Gravity Impact: Sham vs GVS | 1.33 | 1.16 | .24 |
|  | High Gravity Impact: Sham vs GVS | 1.03 | 0.12 | .90 |
| Attempts [Count Variable]<sup>b</sup> |  |  |  |  |
| | Overall interaction | $\chi^2 = 3.04$ | | .22 |
|  | Low Gravity Impact: Sham vs GVS | 0.98 | -0.14 | .88 |
|  | Medium Gravity Impact: Sham vs GVS | 0.87 | -1.87 | .06 |
|  | High Gravity Impact: Sham vs GVS | 1.06 | 0.66 | .51 |
| Time [Continuous Variable] (Log-Linear)<sup>c</sup> |  |  |  |  |
| | Overall interaction | $\chi^2 = 3.55$ | | .17 |
|  | Low Gravity Impact: Sham vs GVS | 1.08 | 1.99 | .047* |
|  | Medium Gravity Impact: Sham vs GVS | 1.05 | 1.44 | .15 |
|  | High Gravity Impact: Sham vs GVS | 0.97 | -0.65 | .51 |
| Distance [Continuous Variable] (Log-Linear)<sup>c</sup> |  |  |  |  |
| | Overall interaction | $\chi^2 = 4.46$ | | .11 |
|  | Low Gravity Impact: Sham vs GVS | 1.11 | 0.80 | .42 |
|  | Medium Gravity Impact: Sham vs GVS | 0.98 | -0.16 | .87 |
|  | High Gravity Impact: Sham vs GVS | 0.77 | -2.13 | .03* |
| Switching [Proportion Variable](Ordered Beta)<sup>d</sup> |  |  |  |  |
| | Overall interaction | $\chi^2 = 2.89$ | | .23 |
|  | Low Gravity Impact: Sham vs GVS | 0.83 | -1.28 | .20 |
|  | Medium Gravity Impact: Sham vs GVS | 1.12 | 1.14 | .25 |
|  | High Gravity Impact: Sham vs GVS | 1.03 | 0.25 | .80 |

*Note.* Overall interactions were evaluated via Wald Type-III Chi-square tests.983 <sup>a</sup>Generalized Linear Mixed Model fit via glmer984 <sup>b</sup> Generalized Linear Mixed Model fit with a zero-truncated negative binomial distribution (glmmTMB).986 <sup>c</sup> Linear Mixed Model (lmer) fit with a natural log-transformed dependent variable.987 <sup>d</sup> Ordered Beta Regression Model fit via glmmTMB.
\*p < .05

At game level, we found a significant negative impact of the GVS in a subset of games only (see Fig. 2). GVS had a negative impact on the success rate in *‘Remove’* (*t*(42) = 2.12, p = .02, Cohen’s *d* = 0.63) and *‘GoalMove’* (*t*(41) = 1.87, p = .03, Cohen’s *d* = 0.56). We also found a detrimental effect of the GVS on the number of attempts for two games, ‘*Spiky’* (*t* (41) = -2.10, p = .02, Cohen’s *d* = -0.63) and *‘Falling_A’* (*t* (42) = -1.78, p = .04, Cohen’s *d* = -0.53). In a majority games, time per attempt negatively correlated with the number of attempts under both stimulations, suggestive of within-game learning. But as no instruction was given on time, these results should be interpretated with caution (see Extended Data Fig. 1).

**Fig. 2.**
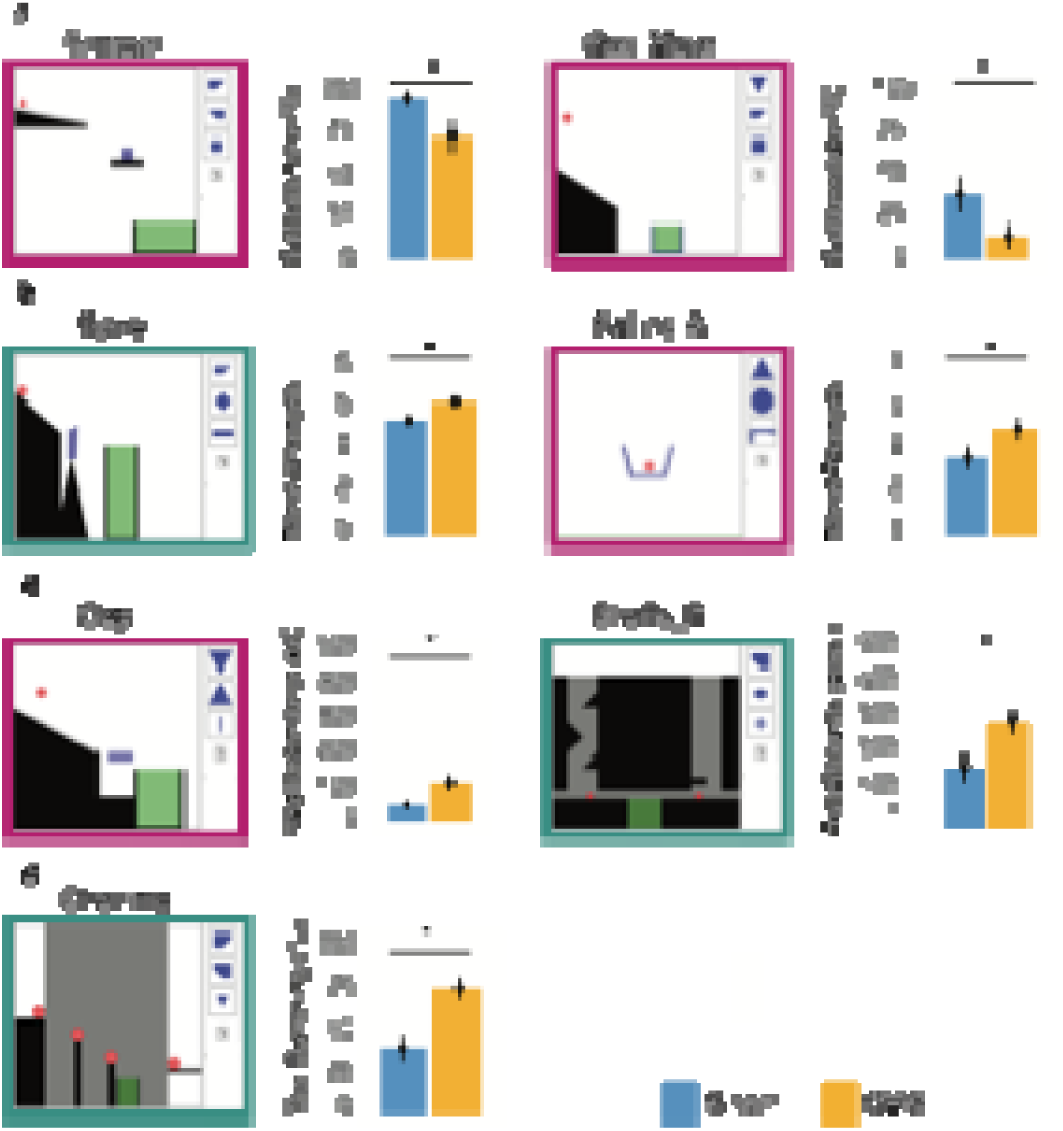
Study 1- Subset of games with a significant impact of the GVS. **a**, on the success rate. The games significantly impacted by the GVS are displayed on the left with their name below the illustration. The bar chart on the right shows the average success rate for the game when played under the Sham stimulation (blue bar) and the GVS stimulation (orange bar). The colour of the borders indicates the impact of gravity in the game. High-gravity-dependent games: turquoise box; Medium-gravity-dependent-games: magenta box. b, on the number of attempts. c, on the distance between placements. d, on the tool switching across attempts. Independent t-tests * *p* < .05

### Study 1: Real-Time Embodied Experience Shapes Reasoning Strategies

We next examined whether the altered vestibular input influenced participants’ high-level strategies - defined as changes in tool selection and spatial positioning across successive attempts within each game - as these patterns reflect how participants adapt their behaviour to solve the task (see Methods). Because these measures were averaged across attempts within a game, persistent large shifts in tool positioning and frequent tool switching provide a proxy for poor evaluation of the failed outcome rather than adaptive exploration, as validated empirically in previous work using the Virtual Tools ^20,32,33^. In their successive placements across games, participants used not-significantly larger distance between their tool placements in the GVS condition compared to the Sham but switched between tools near significantly more in the Sham condition compared to the GVS condition (*t*_distance_(43) = -0.52 *p*_distance_ = .61, Cohen’s *d*_distance_ = -0.11; *t*_tool_switching_(43) = 1.9, *p*_tool_switching_ = .06, Cohen’s *d*_tool_switching_ = 0.28; See full statistics in Table 1). Although the global interaction between stimulation type and gravity impact level did not reach statistical significance for either distance or switching rate (all χ^2^ ≤ 4.46, all *p*s ≥ .11, see full statistics in Table 2 and Methods), pairwise contrasts revealed a significant effect of the stimulation for the high gravity impact games (t_distance_high_(662)= -2.13, p_distance_high_ = .03), with the average distance across attempts 23.2% smaller under Sham than GVS.

As for performance measures, the effects of GVS were specific to a subset of games (see Fig. 2). Participants in the GVS condition placed tools significantly farther apart across attempts compared to Sham in two games, ‘*Shafts_B’* (*t*(25) = -2.29, *p* =.03, Cohen’s *d* = - 0.85) and ‘*Gap*’ (*t* (23) = -2.30, *p* = .03, Cohen’s *d* = -0.90). Regarding tool switching, in ‘*Chaining*’, participants under GVS switched between tools more frequently than under the Sham condition (*t*(36) = -3.27, *p* = .002, Cohen’s *d* = -1.04).

To determine whether the observed differences in distance between successive tool placements reflect distinct reasoning strategies induced by altered gravitational signalling - rather than incidental performance fluctuations, we conducted two complementary exploratory analyses (not pre-registered; see Methods). These analyses were used to characterize the spatial and sequential organisation of tool-use strategies and they explicitly tested if the distances between tool placements across attempts could reliably distinguish stimulation groups, directly quantifying differences in participants’ strategic approaches. In the first analysis, we explicitly investigated whether GVS increased the likelihood of participants changing their spatial tool placement strategies between attempts. Using a Dirichlet Process Gaussian Mixture Model on participants’ placement coordinates, we quantified how frequently participants switched from one identified strategy cluster to another (Fig. 3a). This analysis confirmed that GVS significantly increased the probability of switching strategies compared to Sham stimulation (odds ratio [OR] = 0.83, 95% confidence interval [CI] = [0.70, 0.98], p = .03 - Fig. 3b), and that switching probability decreased over successive attempts (OR = 0.87, 95% CI = [0.83, 0.92], p < .001), suggesting stabilization of strategy use across trials. We then conducted a second complementary analysis using a leave-one-out kernel density classification approach to address whether participants adopted systematically different strategies across stimulation conditions. Here, we assessed if a participant’s overall distance in tool placements patterns could reliably predict their stimulation group (GVS or Sham). Predictive accuracy was computed for each left-out participant by comparing their placement patterns against those of other participants. Because the observed accuracy exceeds the range expected by chance (Fig. 3c), this result confirms that altered vestibular signaling systematically influenced participants’ spatial strategies, leading to reliably distinct patterns between stimulation groups.

**Fig. 3.**
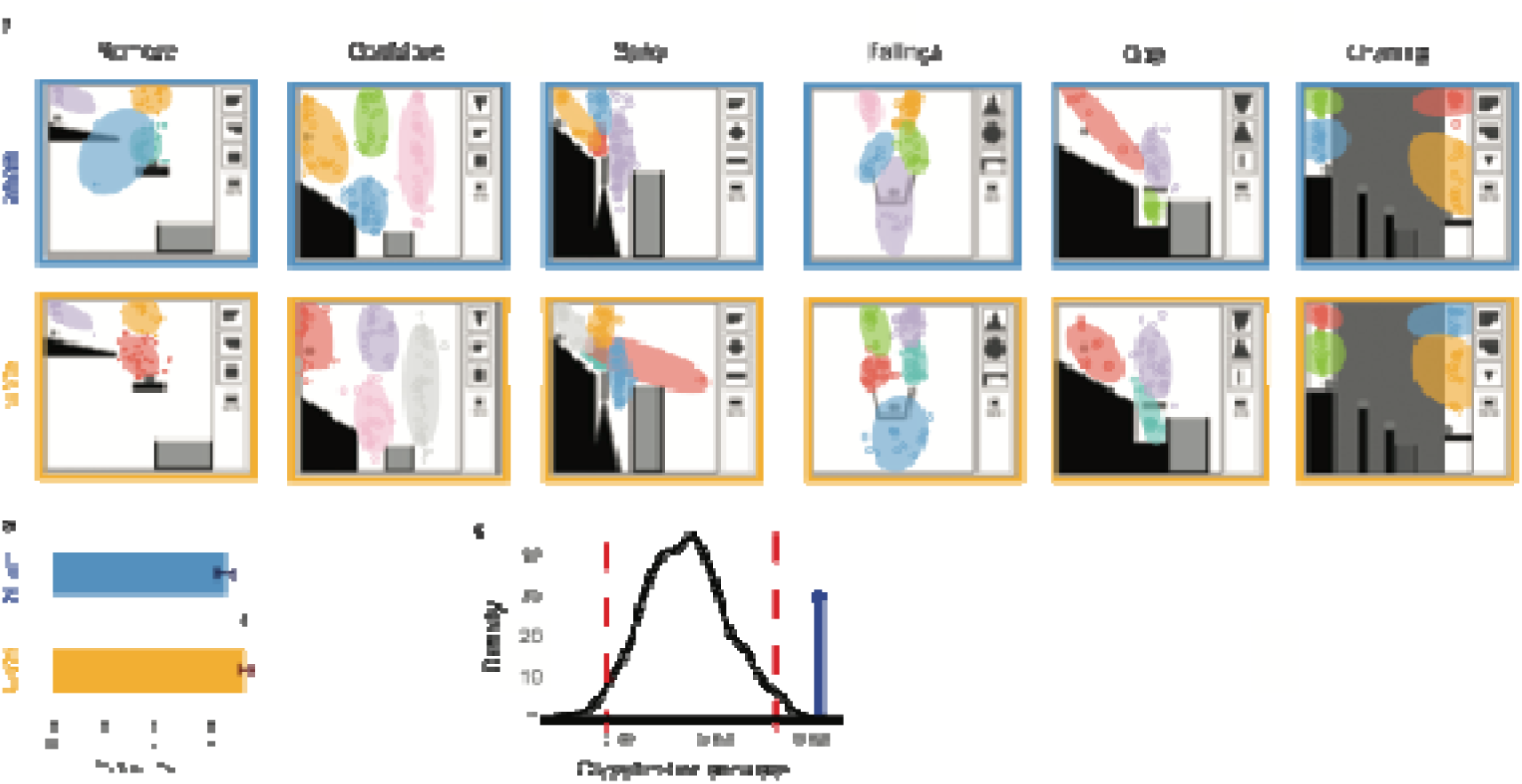
Clustering analysis. **a,** Illustration of results of the Dirichlet Process Gaussian Mixture Model by stimulation type for a selection of games. Each dot represents an individual placement, the ellipses represent the different clusters or strategies b, Strategy switch. Probability of strategy switch on overall games played under GVS versus Sham. c, Leave-one-out Kernel density analysis. This figure shows the distribution of accuracies obtained from 1000 iterations with randomly shuffled group labels and their corresponding 95% Equally-Tailed Intervals (ETIs). The black curve represents the null distribution obtained by randomizing group labels; red dashed lines mark its 95% credible interval. The blue bar marks the observed accuracy, which exceeds this range—indicating that tool placement patterns reliably distinguish between groups.

### Study 2: Real-time embodied experience supports physical reasoning adaptation

In Study 2, participants performed the Virtual Tools games in non-terrestrial gravity. The task required reasoning about hypogravity (0.5 g) or hypergravity (2 g) while experiencing GVS or Sham condition. Similar to previous research manipulating visual gravity in Virtual Tools^34^ and showing no difference in the effect of hypo vs hyper gravity on physical reasoning, we found no difference in the effect of the stimulation on success rates in hypo and hyper gravity games (hypogravity: t(39) = -0.43, *p* = .66, Cohen’s *d* = -0.09; hypergravity: t(39) = 0.10, *p* = .92, Cohen’s *d* = 0.02). Adding the gravity level in the mixed-effects models did not result in this factor being a significant predictor for any of the performance or strategy measures (all *p*s >.05), so we aggregated 0.5 and 2g results as ‘altered gravity’ in subsequent analyses. This aggregation should not be interpretated as evidence that 0.5g and 2g are cognitively equivalent, but rather that no reliable hypo/hyper gravity distinction was detected in the present task and parameter ranges.

Across all games, participants succeeded not significantly better in the GVS condition compared to the Sham condition, with similar number of attempts (*t_success_rate_*(39) = -0.18, *p_success_rate_* = .57, Cohen’s *d_success_rate_* = -0.03; *t_number_of_attempt_*(39) = -0.64, *p_number_of_attempt_* = .26, Cohen’s *d_number_of_attempt_* = -0.09; See full statistics in Table 3). Neither the interactions between gravity impact level and stimulation type nor any of the subsequent pairwise contrasts reached statistical significance (see full statistics in Table 4). At game level, we found a significant negative impact of the GVS in a subset of games only (see Fig. 4). GVS had a negative impact on the success rate in *‘Goal Move’* (*t*(37) = 1.85, *p* = .04, Cohen’s *d* = 0.58). We found a detrimental effect of the GVS on the number of attempts for two games, ‘*Towers_B’* (*t* (38) = -1.96, *p* = .03, Cohen’s *d* = 0.61) and *‘CatapultAlt’* (*t* (37) = -1.78, *p* = .04, Cohen’s *d* = -0.56). We also observed a detrimental effect of GVS on time per attempt, for one game, ‘*Chaining’* (*t*(38) = -1.74, *p* = 0.044, Cohen’s *d* = -0.54). In a majority of games, time per attempt negatively correlated with the number of attempts in both stimulations, suggestive of within-game learning (Extended Data Fig. 2).

**Fig. 4.**
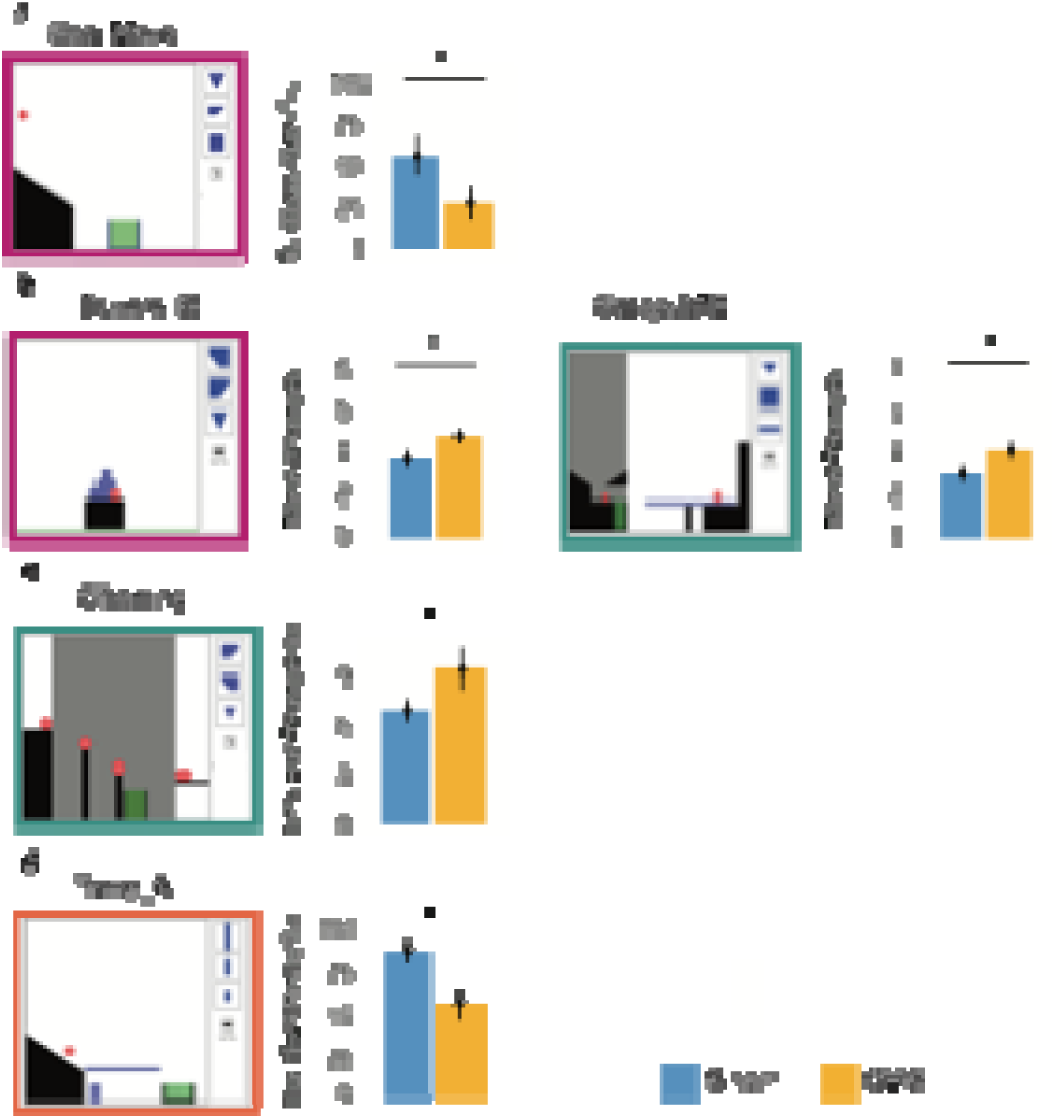
Study 2- Subset of games with a significant impact of the GVS. **a**, on the success rate. The games significantly impacted by the GVS are displayed on the left with their name below the illustration. The bar chart on the right shows the average success rate for the game when played under the Sham stimulation (blue bar) and the GVS stimulation (orange bar). The colour of the borders indicates the impact of gravity in the game. High-gravity-dependent games: turquoise box; Medium-gravity-dependent-games: magenta box; Low-gravity-dependent-games: dark orange box. b, on the number of attempts. c, on the time per attempt. d, on the tool switching across attempts. Independent t-tests, * *p* < .05

**Table 3.** Means, Standard Deviations and t test Statistics for Study 2.

| Variable | Sham |  | GVS |  | df | t | p | Cohen's<br>d |
| --- | --- | --- | --- | --- | --- | --- | --- | --- |
|  | M | SD | M | SD |  |  |  |  |
| Success Rate (%) | 66.7 | 19.6 | 67.3 | 16.8 | 39 | -0.18 | .57 | -0.03 |
| Number of Attempts | 3.2 | 0.9 | 3.3 | 1.0 | 39 | -0.64 | .26 | -0.09 |
| Time per Attempt (s) | 7.7 | 2.5 | 7.7 | 3.0 | 39 | 0.08 | .53 | 0.01 |
| Distance (pixel) | 142.1 | 52.0 | 139.2 | 54.5 | 39 | 0.27 | .78 | 0.05 |
| Tool Switching (%) | 63.7 | 13.7 | 64.1 | 16.7 | 39 | -0.14 | .89 | -0.03 |

Paired t-tests. $p < .05$

**Table 4.**
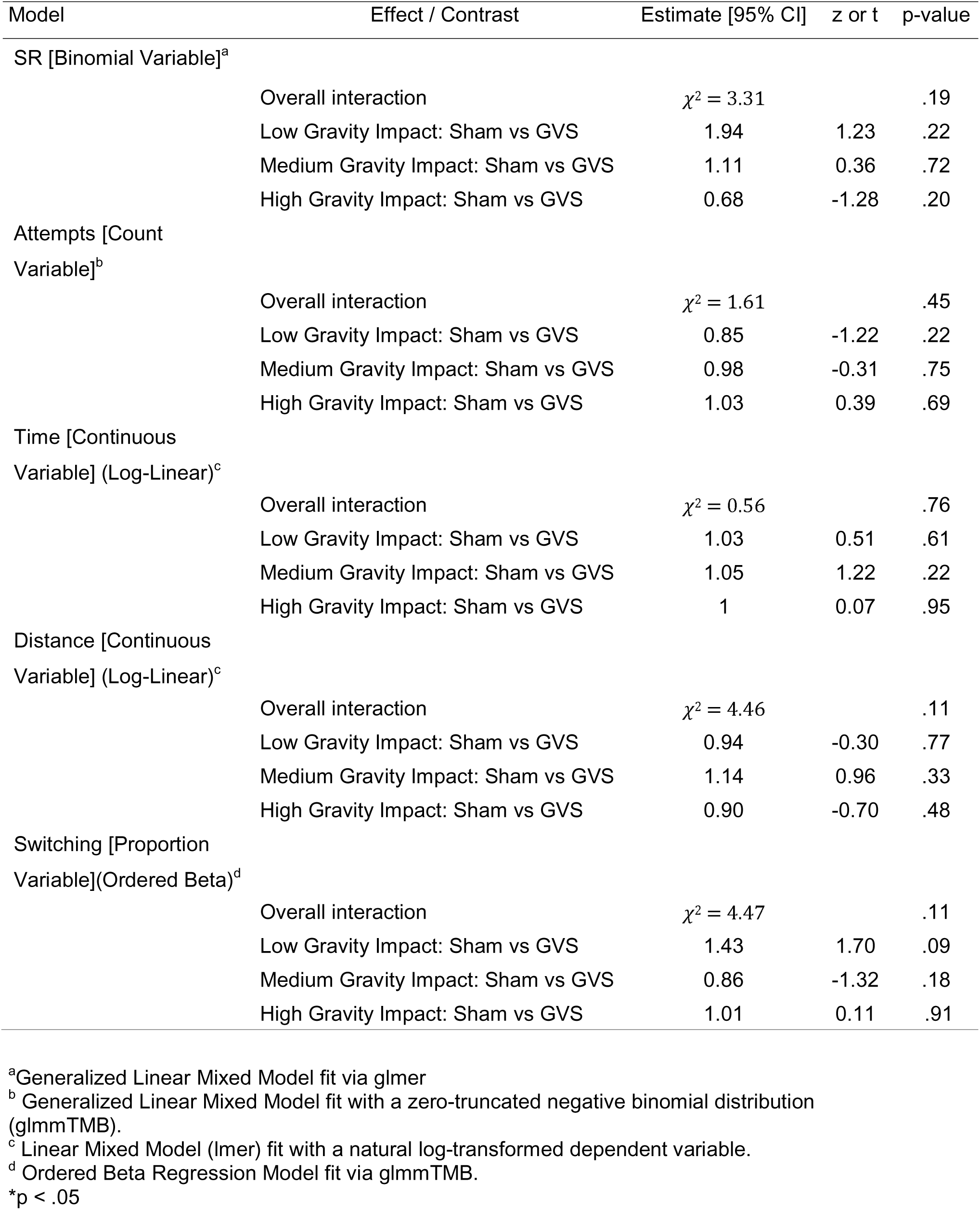
Summary of overall interactions and pairwise contrasts for Study 2.

| Model | Effect / Contrast | Estimate [95% CI] | z or t | p-value |
| --- | --- | --- | --- | --- |
| SR [Binomial Variable]<sup>a</sup> |  |  |  |  |
| | Overall interaction | $\chi^2 = 3.31$ | | .19 |
|  | Low Gravity Impact: Sham vs GVS | 1.94 | 1.23 | .22 |
|  | Medium Gravity Impact: Sham vs GVS | 1.11 | 0.36 | .72 |
|  | High Gravity Impact: Sham vs GVS | 0.68 | -1.28 | .20 |
| Attempts [Count Variable]<sup>b</sup> |  |  |  |  |
| | Overall interaction | $\chi^2 = 1.61$ | | .45 |
|  | Low Gravity Impact: Sham vs GVS | 0.85 | -1.22 | .22 |
|  | Medium Gravity Impact: Sham vs GVS | 0.98 | -0.31 | .75 |
|  | High Gravity Impact: Sham vs GVS | 1.03 | 0.39 | .69 |
| Time [Continuous Variable] (Log-Linear)<sup>c</sup> |  |  |  |  |
| | Overall interaction | $\chi^2 = 0.56$ | | .76 |
|  | Low Gravity Impact: Sham vs GVS | 1.03 | 0.51 | .61 |
|  | Medium Gravity Impact: Sham vs GVS | 1.05 | 1.22 | .22 |
|  | High Gravity Impact: Sham vs GVS | 1 | 0.07 | .95 |
| Distance [Continuous Variable] (Log-Linear)<sup>c</sup> |  |  |  |  |
| | Overall interaction | $\chi^2 = 4.46$ | | .11 |
|  | Low Gravity Impact: Sham vs GVS | 0.94 | -0.30 | .77 |
|  | Medium Gravity Impact: Sham vs GVS | 1.14 | 0.96 | .33 |
|  | High Gravity Impact: Sham vs GVS | 0.90 | -0.70 | .48 |
| Switching [Proportion Variable](Ordered Beta)<sup>d</sup> |  |  |  |  |
| | Overall interaction | $\chi^2 = 4.47$ | | .11 |
|  | Low Gravity Impact: Sham vs GVS | 1.43 | 1.70 | .09 |
|  | Medium Gravity Impact: Sham vs GVS | 0.86 | -1.32 | .18 |
|  | High Gravity Impact: Sham vs GVS | 1.01 | 0.11 | .91 |

<sup>a</sup>Generalized Linear Mixed Model fit via glmer<sup>b</sup> Generalized Linear Mixed Model fit with a zero-truncated negative binomial distribution (glmmTMB).<sup>c</sup> Linear Mixed Model (lmer) fit with a natural log-transformed dependent variable.<sup>d</sup> Ordered Beta Regression Model fit via glmmTMB.
\*p < .05

Across the two studies, there was no significant difference in the average success rates in the baseline games played in both studies (independent two-sided *t*-test: *t*_(82)_ = 0.15, *p* = .88, Cohen’s *d* = 0.03). Potential differences in the GVS effect in altered vs terrestrial gravity could therefore not be explained by potential differences in participants’ reasoning performance between the two studies. To compare the effect of the GVS on physical reasoning in terrestrial vs altered gravities, we defined a Gravity-Index (GI) as the ratio between results for each measure in GVS and Sham, at each gravity impact level (see equation (1) in Methods: this index was not pre-registered). Because of the distribution of these indexes (see Extended Data Fig. 3), we analysed them using Bayesian cumulative link models (see Methods). For all the performance indexes, the interactions between study and gravity impact level were uniformly non-significant across models, with the exception of a marginal trend for time. This model yielded a slight negative interaction estimate for Study 1 at low gravity impact level (posterior median = -0.22, 89% CI [-0.53, 0.09]), indicating a shorter time from the grand mean. Following our hypotheses, we proceeded with contrasts. For success rates (Fig 5A), the posterior median odds ratio indicated a directional trend for the high gravity impact level, with a lower index in Study 1 than in Study 2 (OR = 0.51, CI [0.24, 1.08]), and no meaningful differences in the other levels (see full statistics in Table 5). For attempts (Fig. 5B), no reliable difference between experiments was observed, as all credible intervals widely overlapped the point of group equivalence. Finally for time (Fig. 5C), the results revealed a directional trend for the low gravity impact level, with a lower index score for Study 1 (OR =0.43, CI [0.18, 1.04]), but no reliable difference for the games where gravity matters.

**Fig. 5.**
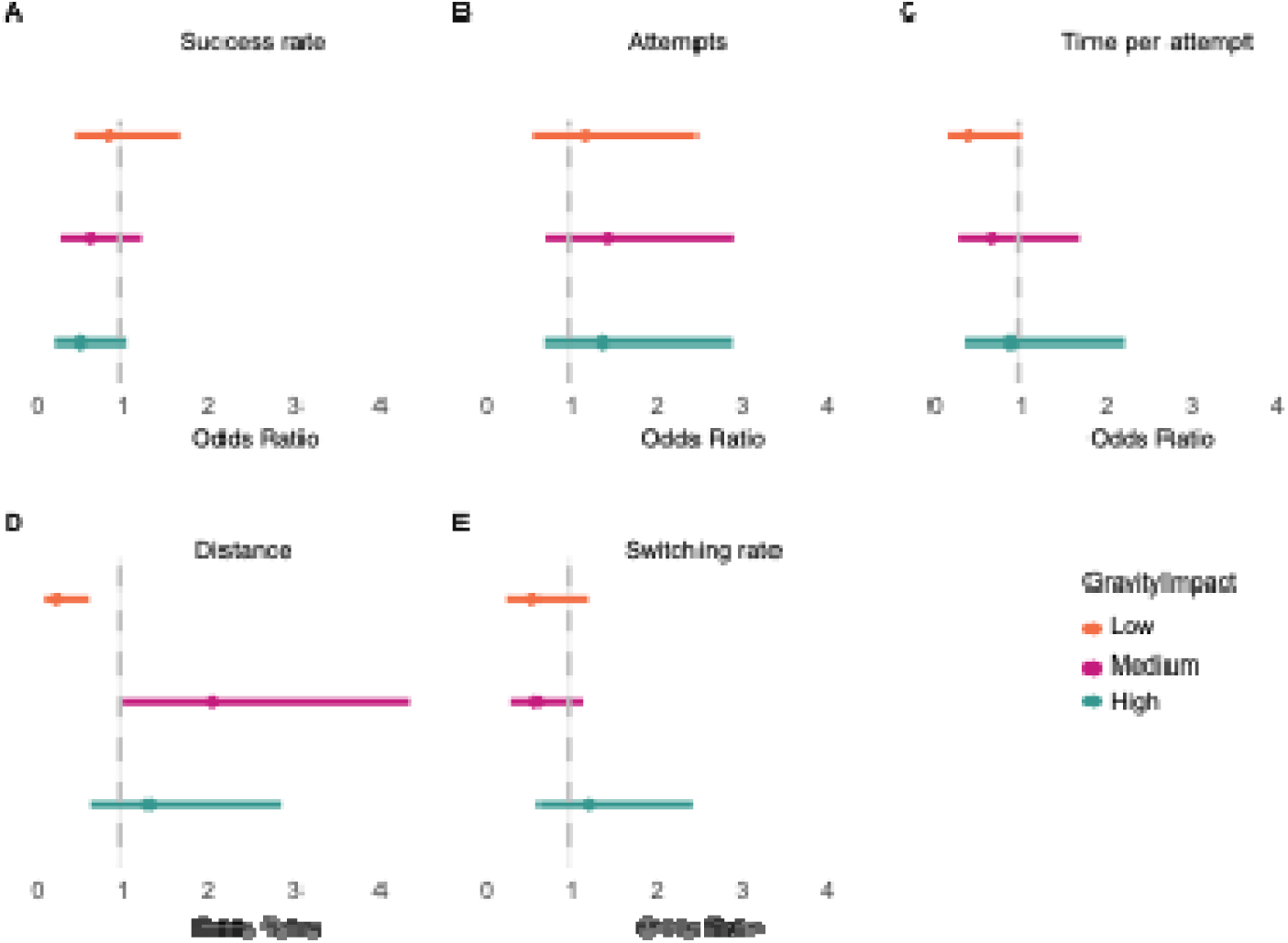
Cross-study Gravity-Index contrast comparisons by gravity impact level. **A,** Success rate. Odds Ratios calculated at 89% Credible Intervals (study 1 – terrestrial gravity – vs study 2 – altered gravity). A contrast is considered to have strong statistical evidence if its 89% CI excludes 1. B, Number of attempts. C, Time per attempt. D, Distance between tool placements. E, Tool Switching between attempts

**Table 5.** Posterior estimates and credible intervals for contrasts.

| Model | Contrast | Estimate (OR) | 89% CI |
| --- | --- | --- | --- |
| SR |  |  |  |
|  | Low Gravity Impact: Study1 vs Study2 | 0.87 | [0.45, 1.70] |
|  | Medium Gravity Impact: Study1 vs Study2 | 0.64 | [0.32, 1.27] |
|  | High Gravity Impact: Study1 vs Study2 | 0.51 | [0.24, 1.08] |
| Attempts |  |  |  |
|  | Low Gravity Impact: Study1 vs Study2 | 1.20 | [ 0.60, 2.51] |
|  | Medium Gravity Impact: Study1 vs Study2 | 1.48 | [0.72, 2.94] |
|  | High Gravity Impact: Study1 vs Study2 | 1.41 | [0.70, 2.92] |
| Time |  |  |  |
|  | Low Gravity Impact: Study1 vs Study2 | 0.43 | [ 0.18, 1.04] |
|  | Medium Gravity Impact: Study1 vs Study2 | 0.72 | [0.30, 1.75] |
|  | High Gravity Impact: Study1 vs Study2 | 0.93 | [0.39, 2.26] |
| Distance |  |  |  |
|  | Low Gravity Impact: Study1 vs Study2 | 0.26 | [ 0.11, 0.62] |
|  | Medium Gravity Impact: Study1 vs Study2 | 2.09 | [1.03, 4.38] |
|  | High Gravity Impact: Study1 vs Study2 | 1.33 | [ 0.64, 2.89] |
| Switching |  |  |  |
|  | Low Gravity Impact: Study1 vs Study2 | 0.59 | [ 0.28, 1.24] |
|  | Medium Gravity Impact: Study1 vs Study2 | 0.62 | [0.33, 1.18] |
|  | High Gravity Impact: Study1 vs Study2 | 1.25 | [0.64, 2.48] |

*Note.* All parameters were estimated using a Bayesian ordinal framework with cumulative logit link functions. Estimates represent the median of the posterior distribution [89% CI]. Contrast estimates and intervals have been back-transformed to the odds ratio scale. A contrast is considered to have strong statistical evidence if its 89% CI excludes 1.

### Study 2: Real-time embodied experience does not impact on strategies

Across all games, participants used not significantly larger distances between placements in the Sham compared to the GVS condition (*t_distance_*(39) = 0.27, *p_distance_* = .78, Cohen’s *d_distance_* = 0.05) and switched between tools at similar levels under the two stimulations (*t_tool_switching_*(39) = -0.14, *p_tool_switching_*= .89, Cohen’s *d_tool_switching_* = -0.03, See Table 3). Regarding the categorisation of games by gravity impact level, none of the interactions between gravity impact level and stimulation type nor the contrasts at each gravity impact level resulted as significant for the strategy measures (see Table 4). At game level, participants switched significantly less between tools in only one game, ‘Table_A’ (*t*(19) = 2.37, *p* = .03, Cohen’s *d* = 1.02- See Fig. 4). No significant negative impact of the GVS was observed for distance across successive placements.

The interactions between study and gravity impact level were non-significant both for distance and switching gravity-indexes. But post-hoc comparisons revealed a distinct crossover interaction pattern for distance indexes (Fig 5D). In the low gravity impact level, the index in Study 2 was significantly higher relative to Study 1 (OR = 0.26, 89%CI [0.11, 0.62]). In contrast, for the medium gravity impact level, the effect reversed: Study 1 demonstrated more than double the odds of achieving higher-tier index values compared to Study 2 (OR = 2.09, 89%CI [1.03, 4.38]). Finally, for the high gravity impact level, no reliable difference between experiments was observed, as the credible interval widely overlapped the point of group equivalence. For switching rate, no reliable difference between experiments was observed, as all credible intervals widely overlapped the point of group equivalence (Fig 5E).

## Discussion

Our study tests a body-dependence in human high-level reasoning. Real-time vestibular perturbation via GVS was found to significantly impair participants’ reasoning about object motion under Earth gravity, yet the same perturbation was different when reasoning in altered-gravity environments. This paradoxical dual effect suggests that high-level cognitive judgments about physics are not fixed computations but are dynamically modulated by the body’s ongoing sensory state. In essence, the brain’s internal “physics engine” appears to be embodied and context-sensitive—robustly tuned to terrestrial gravity under stable conditions, but capable of flexibly reconfiguring its assumptions when sensory cues signal a novel gravitational context.

Our data lend strong support to theories of embodied cognition, which posit that high-level reasoning is deeply rooted in the sensorimotor systems of the body^35^. Rather than viewing physical reasoning as an abstract process operating in isolation, our findings show it to be exquisitely sensitive to the state of the body and its interaction with the environment. The evidence points to a context-dependent effect of GVS. Disrupting the vestibular sense degraded performance in the familiar 1g context, indicating that steady bodily feedback normally scaffolds accurate intuitive physics. Yet the same disruption produced reduced or relatively more favourable effects in altered-gravity context, implying that the brain can rapidly reweight sensory inputs to meet task demands ^36^. This ability to reconfigure cognitive strategies based on bodily context reflects a form of neural plasticity on short timescales. It resonates with evidence that vestibular pathways are not only crucial for balance but also project widely to cognitive centers involved in spatial memory, attention, and executive function ^27,37^. In effect, our “common sense” physical reasoning may emerge from neural computations that evolved for sensorimotor control. For example, humans possess an internalised sense of gravity through lifelong experience, and this embodiment of gravity guides both our actions and intuitions ^38^. The present results show that this embodied knowledge is *not* rigid: when the ground truth changes (as in altered gravity), the mind-body system can adapt by drawing on alternative cues (visual, proprioceptive, or even erratic vestibular signals) to revise its predictions. Such sensorimotor integration ensures that cognition remains grounded and can cope with environmental novelty. More broadly, the context-dependent reversal we observed—where perturbing the body *helps* rather than hinders cognition under changed physics—underscores a core principle of embodied intelligence: mental processes exploit bodily feedback and will adjust their operation to maintain alignment with the external world.

In contrast to spaceflight, parabolic flight or other terrestrial analogues that induce full-body alterations of gravitational input, the present study selectively perturbs vestibular signalling (Study 1) and combined vestibular and visual gravity cues (Study 2), while all other physiological systems remain anchored to a stable 1g environment. Although this may initially appear as a limitation relative to more ecologically valid gravitational transitions, evidence from parabolic flight and related analogue research suggests that fully veridical multisensory feedback is not always necessary to elicit adaptive effects. For example, Kunavar, et al. ^39^demonstrated that applying gravitational compensation to only a segment of the arm can improve motor control in genuinely altered gravitational contexts, despite partial proprioceptive manipulation and multisensory conflict between proprioceptive and vestibular signals. This indicates that targeted or partial sensory perturbations can still engage adaptive recalibration mechanisms. What also appears crucial is the coupling in the present study between GVS and physical reasoning. Rannaud Monany, et al. ^40^showed that mere exposure to altered gravitational environments during parabolic flight is insufficient to update internal models in the absence of active task engagement, such as motor imagery or goal-directed behaviour. Thus, integrating vestibular perturbation with active physical reasoning tasks may be a key determinant of adaptive recalibration, even when gravitational changes are not globally applied.

Our findings also invite interpretation through the lens of predictive coding and internal models of physics. The human brain maintains an implicit model of Earth’s gravity that normally aids perception and action ^38,41^, for example by allowing accurate catching of falling objects. In conventional settings this internal model functions as a reliable prior expectation, integrated with vestibular and other sensory inputs to stabilise our experience of the world ^42^. However, a strong gravitational prior can become a liability in an altered-gravity context, where it produces systematic prediction errors. Our results suggest that adding vestibular “noise” via GVS effectively perturbs or downregulates this prior, forcing the cognitive system to rely more on immediate sensory evidence. In predictive-processing terms, the perturbation injects precisely the kind of prediction errors that prompt the brain to update its beliefs about physical dynamics. Notably, there is independent evidence that introducing stochastic vestibular input can *improve* performance by amplifying information throughout in neural circuits ^43^. Consistent with the principle of stochastic resonance, a moderate level of vestibular perturbation may boost the brain’s sensitivity to new gravitational contingencies rather than being purely disruptive ^43^. Sherman, et al. ^44^showed that noisy GVS induces higher order cognitive benefits in some individuals. While further investigation is required to understand which individuals are sensitive to stochastic resonance, the beneficial role of controlled noise connects both with motor research showing that some level of variability is optimal for adaptive performance as well as computational work ^45–47^. Thus, when gravity is altered, the GVS-induced sensory prediction errors appear to accelerate the revision of the brain’s internal model of gravity, yielding more adaptive reasoning. This perspective aligns with recent views that the cerebellum and vestibular pathways implement forward models to predict gravity’s effects. By momentarily destabilising these predictions, GVS allowed new sensorimotor information to recalibrate cognitive expectations on the fly. In short, an embodied predictive-coding mechanism may underlie the flexible physics reasoning observed here, with vestibular signals serving as a key “teaching signal” when our default assumptions about the physical world no longer hold.

The analyses collapsed across all games did not reveal significant overall effects. This pattern is informative as these analyses combined gravity-dependent problems with problems in which gravity does play little or no role in the solution (Extended Data Table 1). If GVS had produced a general dizziness-related, physiological, attentional or arousal effect, we would expect a broad impairment across the task. Instead, the mixed-effects models show that the effects were tied to gravity-relevant contexts. GVS selectively disrupted physical reasoning in games where terrestrial gravity was essential for identifying the correct solution, while leaving performance on less gravity-dependent games relatively unaffected. Given the distributed nature of the vestibular cortical network and its involvement in a broad range of cognitive functions ^48^, this selectivity argues against a non-specific effect of GVS on arousal, attention, or task engagement. Instead, it suggests a more targeted contribution of vestibular reliability to the representation and use of gravitational knowledge during reasoning. This is consistent with previous evidence demonstrating that GVS perturbs spatial cognitive processes. For example, Dilda et al.DD reported impairments in both short-term spatial memory and egocentric mental transformation following GVS. Our findings suggest that vestibular signals contribute not only to spatial cognition but also to higher-order physical reasoning when successful performance depends on accurate predictions about the effects of gravity.

Importantly, the relationship between GVS effects and the gravity demands of the tasks further supports a gravity-specific interpretation. In study 1, vestibular perturbation was associated with a trend-level increase in the number of attempts and a significant increase in placement distance in gravity-dependent games. In contrast, these effects were not observed in study 2, where participants reasoned within an altered-gravity environment. This pattern is reflected in the comparison of gravity indices across studies, which revealed moderate evidence for effects on success rate and significant effects on distance in gravity-dependent games. Together, these findings suggest that vestibular signals support reasoning in accordance with Earth’s gravitational regularities and that disrupting these signals may, under some circumstances, reduce the influence of terrestrial gravity priors, thereby facilitating adaptation to novel gravitational environments. Nevertheless, the precise computational mechanisms underlying these effects remain to be determined. Future studies could investigate whether the observed impairments are specifically driven by gravity processing or whether they also reflect disruptions in reasoning about other physical principles that are tightly coupled to gravity, such as free fall, momentum, or ballistic trajectories. Additionally, participants were not instructed to prioritise either speed or accuracy, raising the possibility that different individuals adopted different speed–accuracy trade-offs across conditions. Therefore the time index should be interpretated cautiously. Future work could address this limitation by incorporating eye-tracking measures, which would provide a more sensitive index of deliberation, uncertainty, and decision dynamics during physical reasoning.

While Virtual Tools relies on physical reasoning, success in the task is supported by other cognitive processes such as executive function and trial-and-error learning. Importantly, the vestibular perturbation also altered participants’ cognitive strategies, particularly their spatial exploration. Clustering of tool placements showed that participants under GVS shifted more frequently between distinct spatial clusters from one attempt to the next than in the Sham condition, indicating noisier and less stable spatial exploration, in line with evidence that GVS modulates spatial perception and biases distance judgements^31^. By contrast, the near-significant increase in tool switching in the Sham condition may seem at odds with the idea that frequent switching reflects poor evaluation of failed outcomes. Given the relatively small number of attempts per game, however, this pattern is better interpreted as suggesting that GVS dampens cognitive flexibility, consistent with previous reports of its detrimental effects on flexibility^49^. Contrary to the clear pattern observed between Study 1 and Study 2 for distance, with reduced gravity index in Study 2 versus Study 1 for games where gravity matters and the reversed pattern for games with low gravity impact, the interpretation of the comparison of the switching rate indexes is more uncertain and could be investigated further in future research.

When gravity within the game was altered, the same vestibular disruption became functionally beneficial. The trend in the success rate index in gravity-dependent games suggests that vestibular noise facilitated adaptation to the recalibrated gravitational field and helped reduce sensory prediction errors. This rapid improvement, achieved after less than 15 minutes of exposure, is striking given the omnipresence of gravity in everyday life and prior evidence that enhanced performance in novel vestibular environments typically follows much longer pre-adaptation to noisy GVS (around 120 minutes)^36^. The effects on the distance index in altered gravity suggests that vestibular perturbation may also start to reshape strategic exploration—a possibility that future studies should examine more systematically. More extreme gravitational manipulations may reveal effects that are not captured by 0.5g and 2g.

Taken together, our findings highlight the remarkable flexibility of the human cognitive apparatus and open several avenues for deeper inquiry. The ability of a momentary vestibular manipulation to reshape high-level reasoning speaks to a brain that continually recalibrates its internal rules to match the external context, even for fundamental concepts like gravity. This challenges the notion of intuitive physics as a fixed, encapsulated module; instead, it appears as a fluid interplay between memory, expectation, and real-time sensation. An exciting implication is that *cognitive plasticity* extends into domains once thought stable, with potential applications in training and rehabilitation. For instance, in astronautics and extreme environments, strategically perturbing astronauts’ vestibular inputs during training might accelerate the update of internal models for novel gravitational fields, thereby speeding up adaptation ^50,51^. Likewise, therapeutic vestibular stimulation could possibly be used to push the brain toward alternative strategies when habitual sensorimotor predictions contribute to errors or biases in perception.

On the theoretical front, our findings encourage an expansion of frameworks like predictive coding to more fully include embodiment: the brain’s “predictions” are not only about sensory events but are inextricably linked to the physical self in a given world state. This perspective dovetails with emerging trends in artificial intelligence and robotics, where embodied agents—equipped with physical sensors and the capacity for real-time feedback—show more robust problem-solving and generalization than disembodied algorithms ^35^. Just as our study demonstrates that a human’s reasoning can improve by perturbing and thereby enriching the sensory input in unfamiliar settings, an embodied AI might benefit from injecting controlled noise or variability to avoid overfitting to one environmental regime. In summary, the present work not only highlights the adaptability in human reasoning under sensorimotor perturbation, but also provides a conceptual bridge between neuroscience, cognitive science, and embodied AI: it suggests that intelligence, whether biological or artificial, may achieve its highest flexibility when it harnesses the dynamics of an *embodied* predictive model of the world.

## Methods

An analysis plan for this study was pre-registered in Open Science Framework (https://osf.io/8vutf). Data and analysis code have been deposited on GitHub at https://github.com/Physical-Cognition-Lab/Adaptability-in-altered-gravity.

### Participants

#### Study 1

A total of 48 healthy adults with no history of neurological and vestibular disorders took part in the study (M = 25.2 years old, SD = 4.9; range of age: 18.1 to 34.3; 38 females, 45 right-handed as per the Edinburgh Handedness Questionnaire^52^). Four participants were excluded from analyses as they reported dizziness and did not complete the experiment. Therefore, the final sample consisted of 44 participants (M = 25.7 years, SD = 4.8; 34 females). All participants gave their written informed consent. An a priori power analysis was conducted using G*Power version 3. Results indicated the required sample size to achieve 95% power for detecting a medium effect, at a significance criterion of α = .05, was N = 40 for a 2-way ANOVA. Thus, the obtained sample size of N = 44 is adequate to test the study hypothesis. Participants were recruited through the Birkbeck College website and were informed that they could withdraw at any time. The study was granted ethical approval by the Psychology Ethics Committee at Birkbeck, University of London (references #2122009 and #2122028). The participants were either compensated with credits or with 10 pounds.

#### Study 2

This study was covered by the same ethical approval as study 1. Participants to study 2 were recruited through the Birkbeck College website and did not participate to study 1. All participants gave their written informed consent. A total of 41 healthy adults with no history of neurological and vestibular disorders took part in the study (*M* = 24.4 years old, *SD* = 5.1; range of age: 18.5 to 34.9; 30 females, 39 right-handed as per the Edinburgh Handedness Questionnaire^52^). One participant was excluded from analyses as they reported dizziness and did not complete the experiment. Therefore, the final sample consisted of 40 participants (*M* = 24.4 years, *SD* = 5.2; 29 females).

### Design

#### Study 1

To test the impact of altered gravity on participants’ physical reasoning and to account for the high intrasubject variability^53^ while keeping the overall experiment and stimulation times in line with standard practices^31^, we used a within-subject design, where each participant played games with concomitant GVS and a control Sham stimulation. All the games were designed in terrestrial gravity (Fig. 1 and Extended Data Fig. 4). After practicing four games, each participant played twelve baseline games without any stimulation, allowing us to compare the two studies against a baseline. Then, participants completed two sets of fourteen games with concomitant GVS or Sham stimulations. To mitigate for carry-over and practice effects with the games and the stimulations, we counterbalanced the order of the stimulations and the order of sets of games across participants, resulting in four groups. In addition, we randomized the order of the games. Finally, each participant would only play a given game once during the experiment.

#### Study 2

Participants played a version of the same games designed in altered terrestrial gravity (Fig. 1 and Extended Data Fig. 4). After practicing four games, each participant played nine baseline games in terrestrial gravity and without any stimulation. These baseline games were chosen from the baseline games used in Study 1. Then, participants completed two sets of ten games in altered gravity with concomitant GVS or Sham stimulations (counterbalanced across participants). Half of the games were in half-terrestrial gravity and half of them in double-terrestrial gravity (Fig. 1, Extended Data Table 1 and Supplementary Videos 2 and 3). A red text above the game indicated the gravity in the game. The number of games included in Study 2 was slightly reduced compared to Study 1 to a number deemed to be sufficient for assessing the effects of altered gravity on reasoning. The split of the games between the two sets was slightly amended compared to Study 1 to ensure a balanced mix of games in terms of gravity level, difficulty and gravity dependency. As in Study 1, to mitigate for carry-over and practice effects with the games and the stimulations, we counterbalanced the order of the stimulations and the order of sets of games across participants, and we randomized the order of the games. Finally, each participant would only play a given game once during the experiment.

### Experimental procedure

Participants performed the experiment in a dimly lit room, seated, with their head restrained by a chinrest, facing a screen placed approximately at 30 cm and centered at eye level. After participants provided their consent and completed questionnaires assessing their eligibility to receive the GVS and their hand laterality, they received explanations on the Galvanic Vestibular Stimulation. The experimenter then placed the electrodes and let the participants try the GVS for 10 seconds^31^. After participants confirmed they agreed to proceed with the stimulation, they were given standardized instructions on the task. They were informed that for each trial they would be able to try up to 8 attempts and a maximum duration of 1 minute. For Study 2, they were instructed that the gravity condition would vary across the games and that they needed to look at the red sentence at the top of the screen to check in which gravity environment they would play each game. Next, they practiced 4 practice games and then the baseline games, allowing them to get familiarized with the Virtual Tools games before the GVS stimulation. Finally, they played 2 sets of games with concurrent stimulation (either GVS or Sham). They moved to the next game either when they found the solution or after 8 failed attempts or after 1 minute had passed. They received visually presented feedback (green tick or red cross) after each attempt. After the last game, the experimenter removed the electrodes and debriefed the participants.

### Physical reasoning task

#### Virtual Tools Games

Participants practiced an adaptation of the ‘Virtual Tools’ task, an online gaming framework developed by Allen, et al. ^20^ which has been used in previous research to assess physical reasoning in adults and children^20,33,34,54^. This task consists of a series of computerized games in which participants are asked to select one of three tools in one click and to place it on the screen in another click, to achieve a goal – getting a red object into a green goal area. Their tool placement triggers physical cascades, which approximate the physics of the real word, including gravity and object-to-object interactions (see Figure 1a and Supplementary Video 1). To succeed, participants must use their knowledge about physical laws such as gravity and collision forces and predict environmental changes and object-to-object interactions, without performing physical motor movement. For each game, participants could attempt to place tools up to 8 times with a time limit of one minute. The game is reset to its initial state after a failed attempt.

#### Block types and gravity manipulation

Participants were exposed to different gravities. In the baseline block, participants played the games designed under terrestrial gravity and no stimulation. Then they played two sets of games with a stimulation on, either GVS or Sham. These sets of games were either designed under terrestrial gravity - Study 1-or altered gravity - Study 2 (See Extended data Figure 4 for details of the games in each study). The games were of different levels of difficulty and varied systematically in the extent to which gravity-related predictions are required for solution (See Extended data Table 1), based on piloting and previous computational modelling work in the lab classifying games’ gravity impact in three levels (low-gravity-dependent, medium-gravity-dependent, high-gravity-dependent). This design is central to the logic of the study: if GVS produced a general disruption, effects should appear broadly across the task, if it affected gravity-dependent reasoning, effects should be most evident in games with higher gravity impact. In each study, the two experimental sets were matched on both gravity impact and difficulty factors. In addition, for Study 2, the sets were matched on the two gravity levels (hyper/hypo gravity).

### Galvanic Vestibular Stimulation (GVS)

Vestibular stimulation was delivered using a commercial stimulator (Good Vibrations Engineering Ltd., Nobleton, Ontario, Canada), controlled via LabVIEW. Carbon rubber electrodes (16 cm²) coated with electrode gel were placed bilaterally over the mastoid processes and secured with adhesive tape. Prior to electrode placement, the application sites were cleaned, and electrode gel was applied to minimize impedance. The stimulation protocol used a stochastic waveform, consisting of an alternating sum-of-sines voltage with dominant frequencies at 0.16, 0.32, 0.43, and 0.61 Hz. This stochastic GVS induces a sense of postural instability without producing consistent or directional illusory motion. To prevent compensatory effects from the non-stimulated side, stimulation was delivered binaurally. A Sham stimulation condition was also implemented, which produced similar skin sensations under the electrodes as the GVS, to control for non-vestibular specific effects. The Sham electrodes were placed on the left and right sides of the neck, approximately 5 cm below the GVS electrodes, which were positioned above the mastoid bones. The maximum intensity for both stimulation conditions was set at 1 mA. These parameters, similar to the ones used in previous studies^31^, were selected to optimize vestibular disruption^55,56^, eliciting a mild sensation of dizziness that dissipated immediately after stimulation^57^. These settings have been shown to mimic spatial disorientation, and suprathreshold GVS is considered an analog to spaceflight. In this experiment, the maximum duration of GVS was 14 minutes (14 games × 1-minute maximum), which has been shown to be well-tolerated, with no lasting effects beyond the stimulation. The cross-study design allowed to maintain the total GVS exposure per participant below 20 minutes, in line with safety guidelines with the GVS^58^.

### Data and statistical analyses

Data and statistical analyses were carried out with MATLAB R2020a (MathWorks), the R software environment (version 4.3.3) for statistical computing and graphics and Python (Jupyter Notebook version 6.4.1) for clustering analyses.

Practice games were not included in the analysis. The completion of a minimum of 10 games under each stimulation was set as a requirement for participants’ data to be included in the analysis. This threshold had been defined to ensure the data were sufficient to assess the effects of the GVS versus Sham stimulations. For each participant and each game, we quantified the success rate, the number of attempts needed and the time per attempt as performance measures. We then quantified the tool switching between attempts and the distance between tool positioning in successive attempts as strategy measures.

For each study, for our three performance measures, we conducted one-sided independent t-tests for game level comparisons under the GVS and Sham stimulations and one-sided paired t-tests for comparing participants’ performance under each stimulation. For strategy measures, we used two-sided t-tests as we had no direction in our hypotheses. Each game was treated as a distinct physical-reasoning problem, and the t-tests were used to localize the games in which the stimulation effect was expressed, not to compare games with one another or to draw broad conclusions from isolated contrasts. Therefore, in this case, no multiple-comparison corrections are required.

For investigating the impact of the stimulations at each gravity impact level, we conducted mixed-effect models tailored to the distribution of the dependent variables. Two outlier games, ‘Spiky’ and ‘SeeSaw’, were excluded from these analyses. We used gravity impact level and stimulation type as fixed factors and participant and game as random effects. The categorical predictors were sum-coded prior to analysis. For the performance measures, we used a logistic regression for success, a zero-truncated negative binomial model for number of attempts and a linear regression applied on a log-transformed variable for time. For the strategy measures, we conducted an ordered beta regression model for the switching rate, and a linear regression on log transformed distance. Pairwise contrasts between stimulation type were restricted to a single a priori comparison within each gravity impact level. Consequently, no multiple-comparison adjustment was required. For Study 2, we tested whether hypogravity and hypergravity should be analysed separately by adding gravity level (0.5g vs 2g) to the mixed-effects models. Gravity level was not a significant predictor for any performance or strategy measure, and this was consistent with previous Virtual Tools work showing no reliable difference between hypo- and hypergravity effects. On this basis, we aggregated the two conditions as “altered gravity” for the main analyses. For all the analyses, the level of significance was always set at *p* < .05.

Finally, to compare the Gravity Indexes between Study 1 and Study 2, we used mixed-effect models that include study and gravity-impact level as fixed factors and participant as random factor. Because the indexes were strictly bounded and showed clustering at structural values (see Extended data Fig. 3 for the distribution of each GI), we analysed them using Bayesian cumulative link models, which are appropriate for bounded, non-normally distributed outcomes^59–62^. We examined the pairwise contrasts using the model’s posterior distribution, with 89% quantile-based credible intervals (CI) to evaluate the proportional odds of scoring differently on the ranked continuous index for each gravity impact level between the two studies.

### Computation of primary measures

#### Performance analysis

For each participant, for each trial and for each attempt, the following data were recorded: success (yes/no), time elapsed (in seconds) between the start of the attempt and the placement of the tool, tool selected and its x,y coordinates on the screen. The data were analysed in MATLAB, by averaging data across games and per stimulation, and by assessing the average number of attempts, success rates and time per attempt under each stimulation.

#### Strategy analysis

For each game and for each stimulation type, we evaluated how participants attempted to solve each game by evaluating two measures: the percent of tool switching and the average tool-positioning distance. We focused only on games in which participants failed to solve the game on the first attempt. To calculate the percent of tool switching, for each game and each attempt (starting from the second attempt), we compared whether the selected tool was identical to the one used in the previous attempt. We evaluated the percent of attempts for each game in which the selected tool was different. We then averaged the percent of tool switching across all games within each stimulation type. For the average tool-use positioning distance, we calculated for each game the average Euclidean distance in pixels between the positioning of the tool in each attempt and the previous attempt. For each game, we averaged this Euclidean distance across all attempts (starting from the second attempt), yielding an average tool-use positioning distance per game. Then, we averaged the positioning distance across all games within each stimulation.

### Gravity-Index

Participants completed the virtual tool-use task in both GVS and sham conditions, allowing us to assess the effects of altered gravity on their performance and strategies. As the games relied on gravity to different extent, we calculated a Gravity-Index (GI) for each measure, defined as the ratio between participants’ performance and strategy outcomes in the GVS versus Sham conditions, at each gravity impact level. This GI is a direct quantification of the effects of real-time vestibular noise on high-level reasoning. We then compared the GI across tasks played under terrestrial and altered gravities.

As an example, let Gravity-Index_SRlow_(respectively Gravity-Index_SRmedium_ and Gravity-Index_SRhigh_) be the averaged index for success rates for low-gravity-dependent games (respectively medium-gravity-dependent games, high-dependent-gravity games) for a participant.

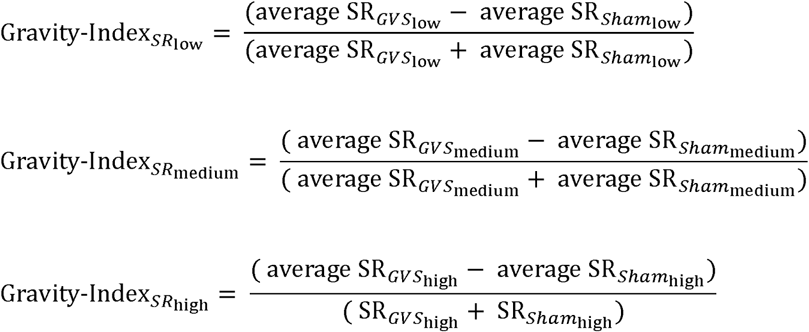

For success GI, a value close to -1 indicates a large negative difference between GVS and Sham gravity results, a value close to 0 similar results and a value close to 1 indicates a large positive difference between GVS and Sham gravity results.

A GI is similarly calculated for each performance and strategy measures for each participant of Study 1 and Study 2, for each gravity impact level. The indexes were calculated in the same way in both studies, and the cross-study comparison was based on games that were common to Study 1 and Study 2, excluding the two outliers games (‘*Spiky*’ and ‘*SeeSaw*’). Then for each performance and strategy measure, we averaged the gravity indexes across participants to calculate the gravity indexes for Study 1 (terrestrial gravity games) and for Study 2 (altered gravity games).

### Dirichlet Process Gaussian Mixture Model

The distance and tool switching measures reduce each participant’s behaviour to two averages; they therefore do not capture the multivariate structure of spatial placements, whether participants move between distinct placement strategies across successive attempts, or whether the overall placement distributions differ systematically between stimulation conditions. To quantify strategic switching behaviour, we employed a Dirichlet Process Gaussian Mixture Model approach adapted from Allen, et al. ^20^. The Dirichlet Process Gaussian Mixture Model is well suited to characterize the spatial and sequential organisation of tool-use strategies because it identifies strategy clusters directly from x/y placement coordinates without imposing an a priori number of strategies, allowing to quantify transitions between spatial strategies across attempts (in relation to the environment in the game which is not taken into account in the more basic measures). For each combination of stimulation condition and trial level, we fitted separate mixture models that captured the unique cluster structure and estimated the probability of each attempt belonging to each identified strategy.

Strategic switches were defined as transitions between different spatial strategies on consecutive attempts within the same game. A switch occurred when the cluster differed between two sequential attempts by the same participant. Given the probabilistic nature of the Dirichlet Process, we repeated the entire clustering procedure across 1,000 iterations.

For each iteration, we fitted a logistic regression model with switch probability as the dependent variable and both stimulation type (GVS vs. Sham) and attempt number as predictors, extracting the p-value for the stimulation type effect. From this bootstrap analysis, we identified the iteration that produced results closest to the modal p-value distribution by creating kernel density estimates of these stimulation effect p-values across all 1,000 iterations. We selected the iteration whose p-value was nearest to the peak density value, representing the most probable statistical outcome under our clustering approach. We report the results from this selected iteration and its corresponding logistic regression model.

### Leave-one-out Kernel density analysis

While the Dirichlet Process Gaussian Mixture Model enables to assess strategic switching behaviour under the GVS and Sham stimulations, it does not address whether participants adopted systematically different strategies across stimulation conditions. To determine whether the placement patterns contain sufficient condition-specific structure to classify stimulation condition above chance, we implemented a leave-one-out kernel density classification approach based on the [x,y] coordinates of each attempt. For each participant, we excluded their data and used the remaining participants’ placements to construct two separate two-dimensional kernel density estimates (KDEs): one for the GVS group and one for the Sham group. These KDEs captured the spatial distribution of tool placements within each condition.

Next, we estimated the probability density of the left-out participant’s placements under each of the two KDEs. These estimates were used to compute a relative likelihood, reflecting how much more likely a given placement was under the correct group distribution compared to the incorrect one. Relative likelihood values were binarized into classification accuracy scores: a value of 1 was assigned if the placement was more likely under the correct group, and 0 otherwise. Accuracy scores were then averaged across all trials for each participant to produce a single accuracy value.

To evaluate whether the observed classification performance exceeded chance, we repeated the procedure for a null model in which stimulation labels were randomly shuffled. This control analysis was run over 1,000 iterations, each time generating KDEs from the permuted data and computing classification accuracy in the same leave-one-out manner. The resulting distribution of accuracies served as a baseline against which the observed accuracy was compared.

Two versions of this analysis were conducted. The first focused exclusively on the first attempt of each game, evaluating early strategy use under each condition. The second was performed at the game level, averaging placement patterns across trials.

## Acknowledgements

This work was supported by the ESRC New Investigator grant ES/W009242/1, BA Talent Award TDA21\210038, Waterloo Foundation grant 917-4975, Leverhulme Trust research grant RPG-2022-327, and the Birkbeck/Wellcome Trust Institutional Strategic Support Fund to OO and by the UK Experimental Psychology Society and BIAL Foundation grant (grant number 041/2020) to E.R.F. We thank Ruben Zamora for helping with the online data collection.

## Author contributions

HG, ERF, and OO conceptualized the study and designed the experiment. HG collected the data. HG and TG analyzed the data under the supervision of ERF and OO. All authors wrote the manuscript and approved the final version of the manuscript for submission.

## Declaration of interests

The authors declare no competing interests.

**Extended Data Fig. 1.**
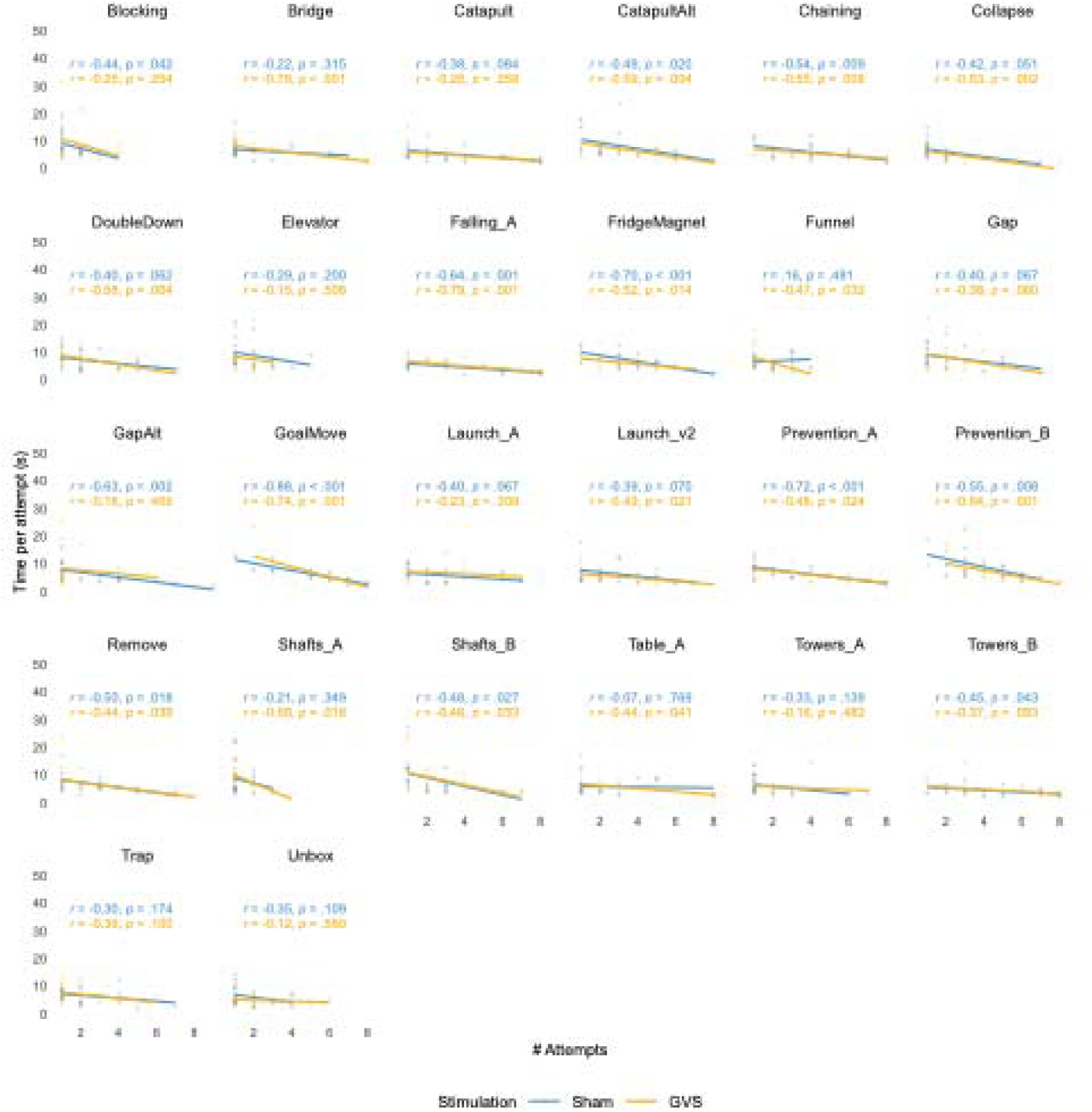
Study 1. Correlation between the number of attempts used and time per attempt for each game played under Sham (blue) and GVS (orange). Dots indicate individual results

**Extended Data Fig. 2.**
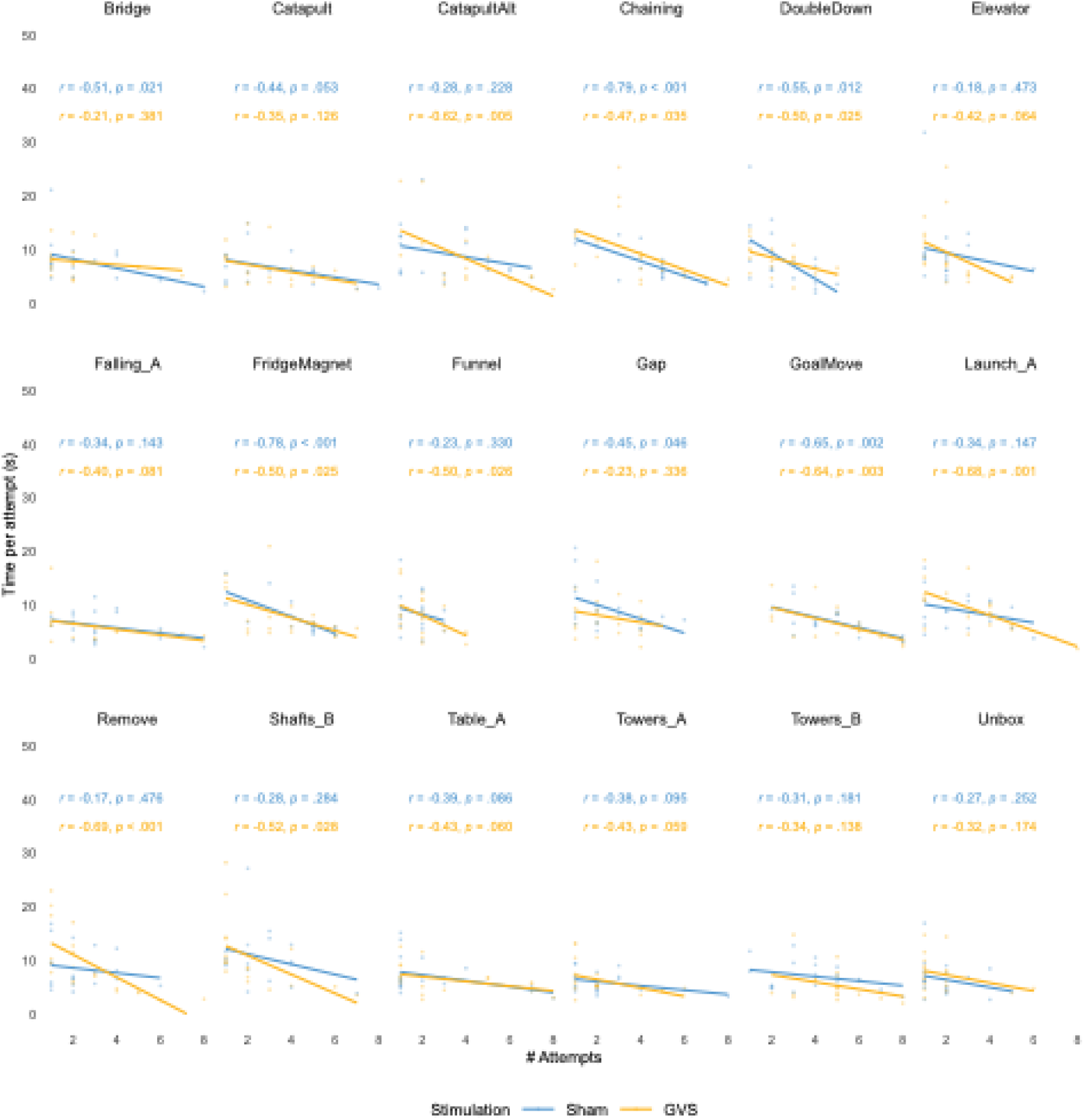
Study 2. Correlation between the number of attempts used and time per attempt for each game played under Sham (blue) and GVS (orange). Dots indicate individual results

**Extended Data Fig. 3.**
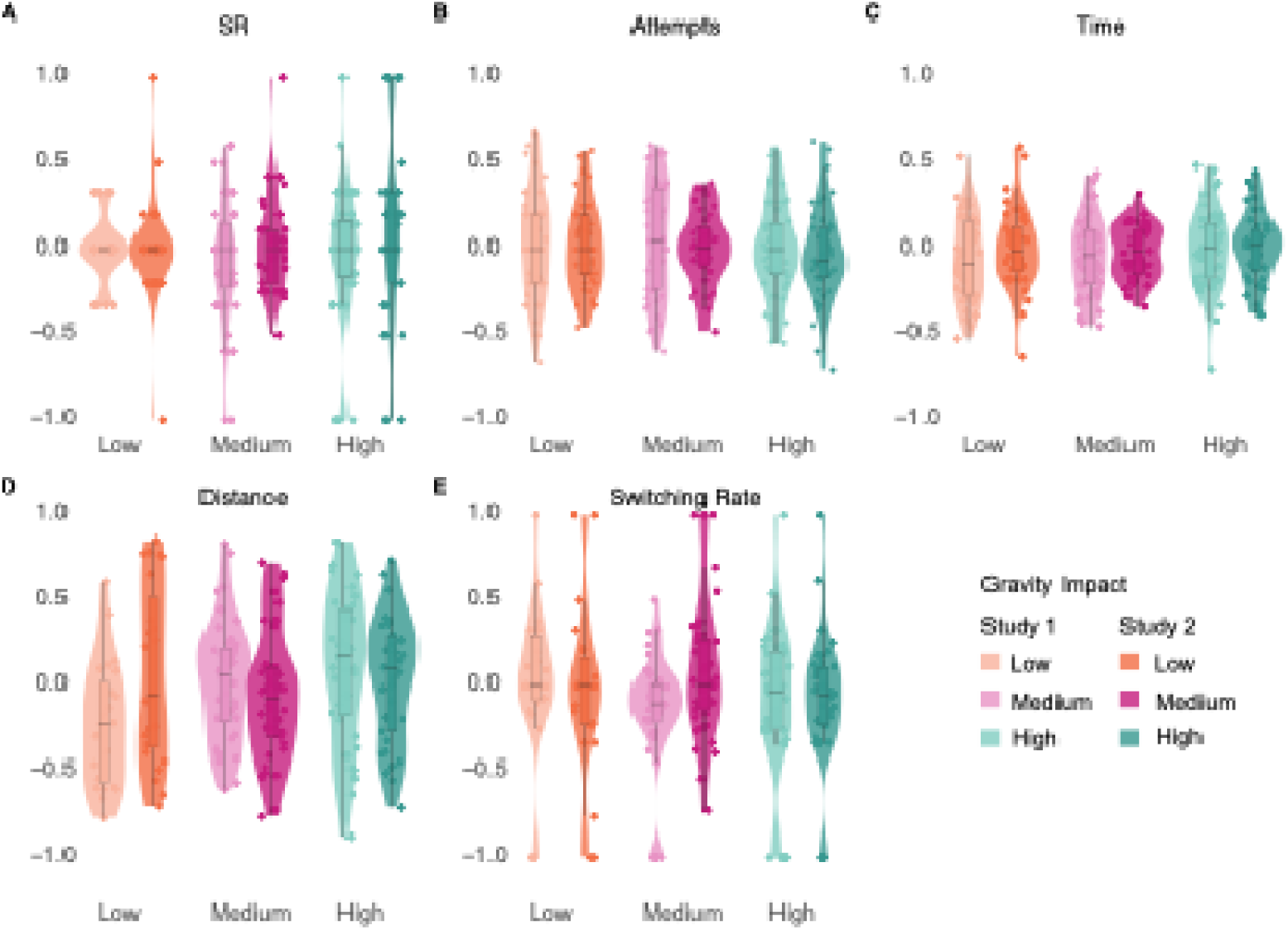
Distribution of Gravity-Index for each measure by study and gravity impact. **A,** Success rate. Each dot represents an individual participant average index for each gravity impact level. Violin boxplot displaying the density of the data, the box plot the median, quartiles and whiskers. For each gravity impact level, the plot on the left represents the Gravity-Index distribution in study 1 (terrestrial gravity) and the plot on the right in study 2 (altered gravity). B, Attempts. C, Time. D, Distance. E, Switching rate

**Extended Data Fig. 4.**
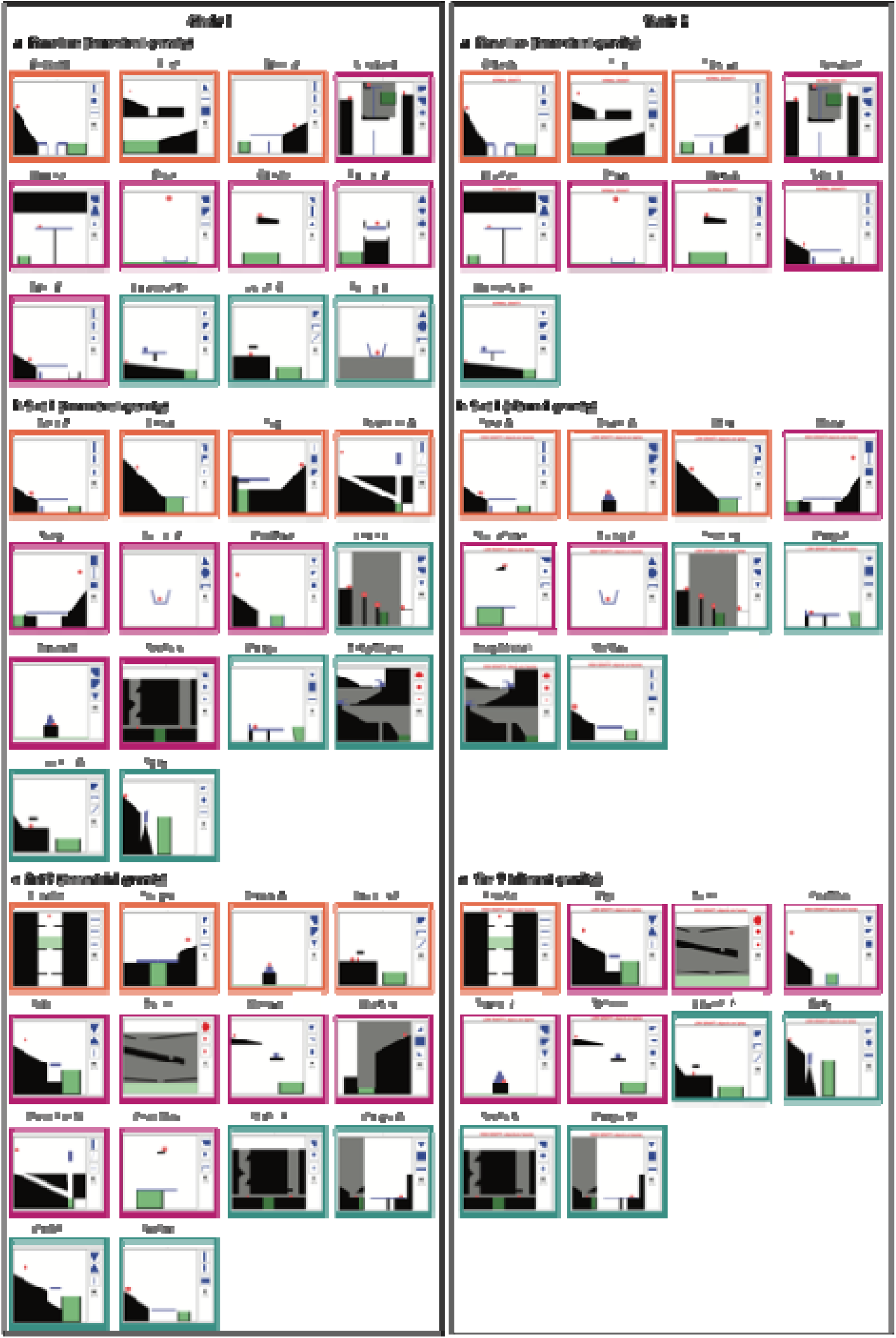
Left panel Games in Study 1. a, Baseline games. b, Games in Set 1. c, Games in Set 2. The colour of the borders indicates the impact of gravity in the game. High-gravity-dependent games: turquoise box; Medium-gravity-dependent-games: magenta box; Low-gravity-dependent games: orange box. Right panel Games in Study 2

**Extended Data Table 1.**
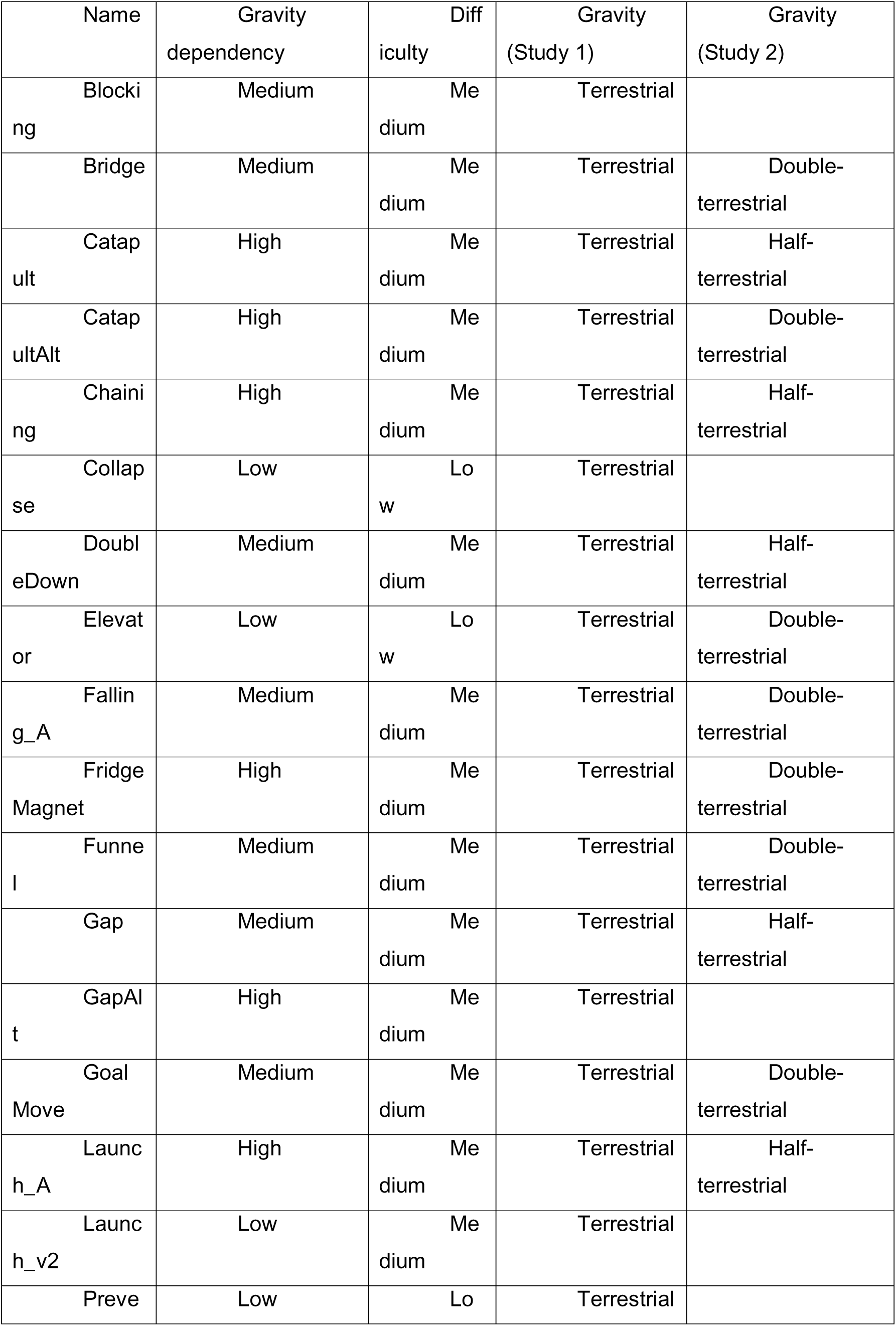
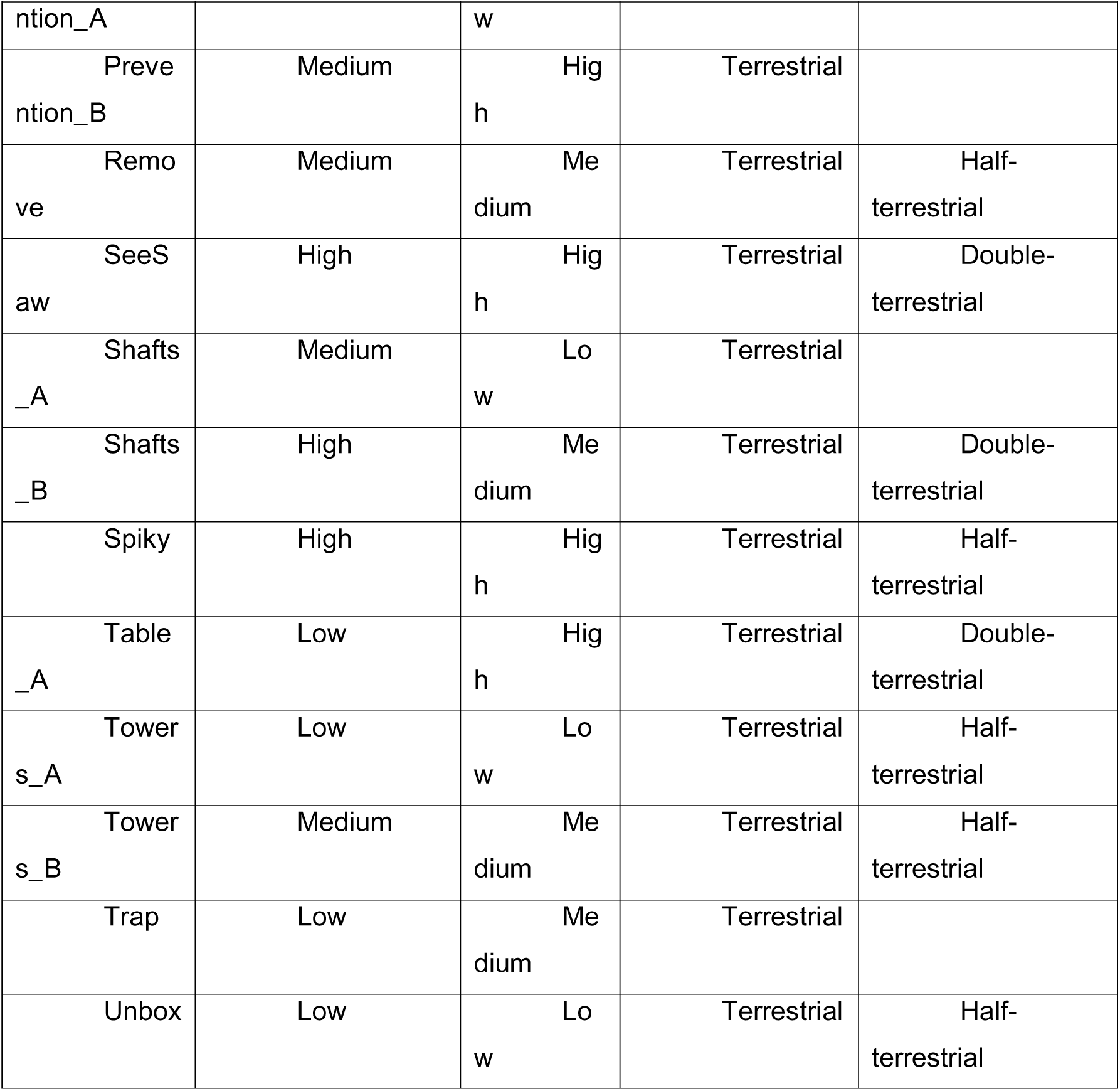
Details of the games played under concomitant stimulation in study 1 and study 2. The gravity dependency column indicates the level of reliance on gravity to solve each game. The difficulty column indicates the level of difficulty to solve the game. The two columns on the right indicate the gravitational accelerations used in each of the two study for a given game.

**Supplementary information Video. 1 Illustrative attempt of a game in terrestrial gravity (1g) – Study 1.** The participant correctly positions the chosen tool on the screen, enabling the red ball to get into the green target area (successful attempt)

File: Catapult_1g

Supplementary information Video. 2 Illustrative attempt of a game in hyper gravity (2g) – Study 2

File: CatapultAlt_2g

Supplementary information Video. 3 Illustrative attempt of a game in hypo gravity (0.5g) – Study 2

File: Catapult_0.5g

